# Liquid droplet germ granules require assembly and localized regulators for mRNA repression

**DOI:** 10.1101/382838

**Authors:** Scott Takeo Aoki, Tina R Lynch, Sarah L Crittenden, Craig A Bingman, Marvin Wickens, Judith Kimble

## Abstract

Cytoplasmic RNA-protein (RNP) granules have diverse biophysical properties, from liquid to solid, and play enigmatic roles in RNA metabolism. Nematode P-granules are paradigmatic liquid droplet granules and central to germ cell development. Here we analyze a key P-granule scaffolding protein, called PGL, to investigate the functional relationship between P-granule assembly and function. Using a protein-RNA tethering assay, we find that reporter mRNA expression is repressed when recruited to PGL granules. We determine the crystal structure of the PGL N-terminal region to 1.5 Å, discover its dimerization and identify key residues at the dimer interface. *In vivo* mutations of those interface residues prevent P-granule assembly, de-repress PGL-tethered mRNA and reduce fertility. Therefore, PGL dimerization lies at the heart of both P-granule assembly and function. Finally, we identify the P-granule-associated Argonaute WAGO-1 as crucial for repression of PGL-tethered mRNA. We conclude that P-granule function requires both assembly and localized regulators.

## Introduction

RNA-protein (RNP) granules, otherwise known as biomolecular condensates ^1^, are ubiquitous non-membrane bound “organelles”. Some RNP granules exist in a solid-like state with little component exchange ^2^, while others behave as liquid-like droplets with components dynamically diffusing in and out of the granule ^3, 4^. Intriguingly, mRNA regulators localizing to liquid granules do not rely on granule association for their activities ^5^. Therefore, despite intense interest and an ever-expanding literature, the functional relationship between granule assembly and mRNA regulation remains a mystery.

Here, we investigate the functional relationship between RNP granule assembly and function in *C. elegans* germline P-granules. P-granules are paradigmatic RNP granules ^6, 7^ with striking similarities to germ granules in *Drosophila* and vertebrates, both in subcellular location and composition ^8^. They are crucial for fertility and totipotency ^9^, and some of their components can display liquid droplet behavior ^6, 10, 11^. In adult germ cells, P-granules localize to the cytoplasmic face of nuclear pores (**Figure 1A**) and contain both untranslated mRNAs ^12^ and numerous RNA regulatory proteins ^13^. Yet their molecular function is poorly defined. RNAi-mediated gene repression is seated there ^14^ and loss of P-granules correlates with aberrant upregulation of somatic transcripts ^15, 16, 17^. Yet it remains unclear whether mRNA repression depends on granule scaffold assembly or on localized regulators in granules. A definitive understanding of this fundamental question requires manipulation of mRNA localization as well as manipulation of granule formation without eliminating pivotal assembly proteins. The former is made possible with a protein-mRNA tethering assay ^18, 19, 20^, but the latter requires a deeper understanding of P-granule scaffolding proteins and the molecular basis of their assembly.

**Figure 1.**
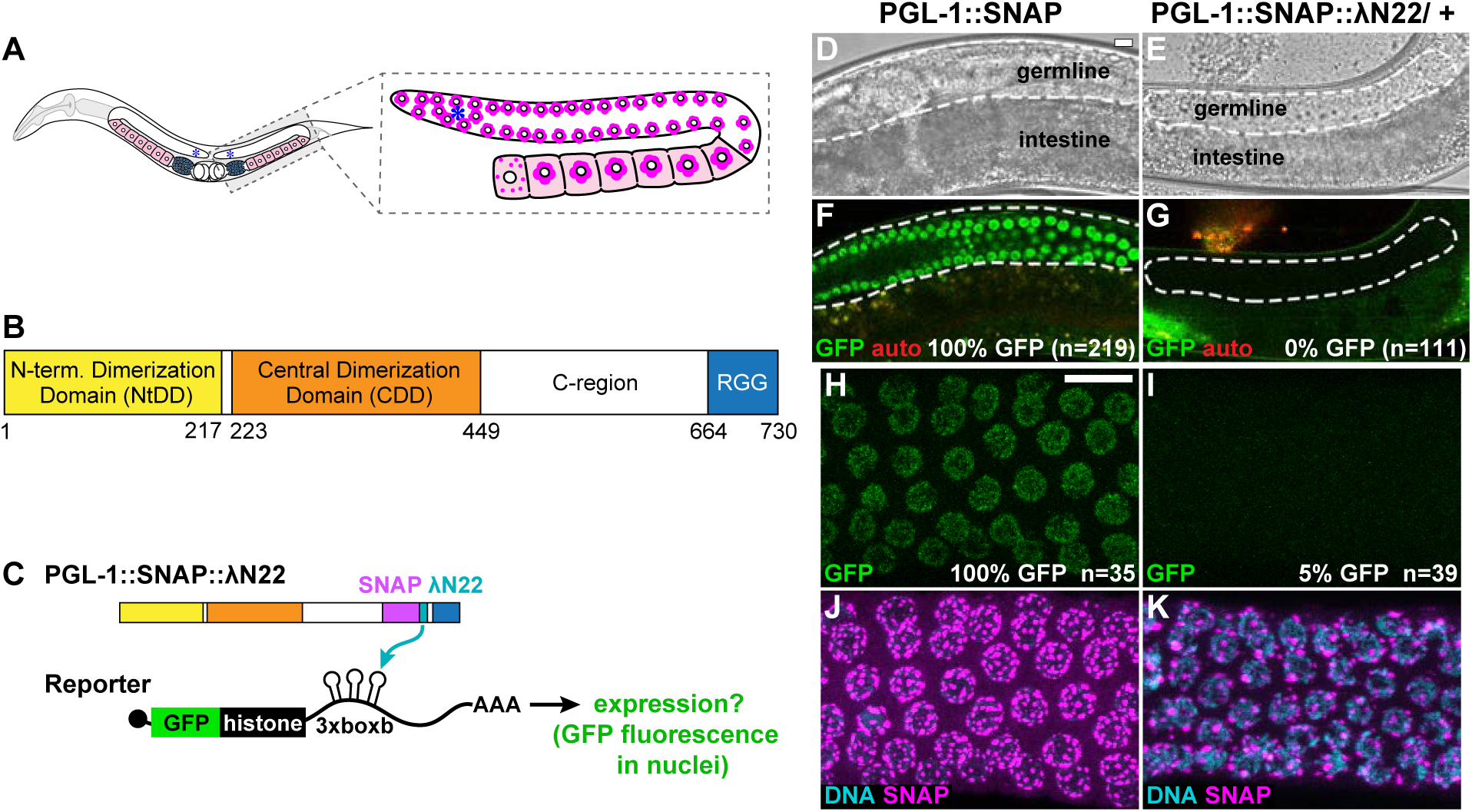
PGL-tethering represses an mRNA reporter *in vivo* (A) Left, *C. elegans* adult hermaphrodite possesses two gonadal arms with proliferating germ cells at one end (asterisk) and differentiating gametes at the other. Gonads make sperm (blue) first and then oocytes (pink). Right, P-granules (magenta) reside at the nuclear periphery of all germ cells until late oogenesis. (B) Linear diagram of *C. elegans* PGL-1. (C) Protein-mRNA tethering assay. The reporter mRNA encodes GFP-histone H2B and harbors three boxb hairpins in its 3’UTR; a ubiquitous germline promoter drives expression (see Methods). λN22 peptide (light blue) is inserted into PGL-1 with a SNAP tag (magenta). Binding of PGL-1::SNAP::λN22 to boxB hairpins recruits reporter mRNA. (D-G) GFP reporter expression in germ cells of live animals. (D,E) Brightfield image. (F,G) GFP fluorescence (green); auto fluorescence (red). n, number of animals scored for GFP expression. “%”, germlines with detectable GFP. Scale bar, 10 μm, in D applies to D-G. (H-K) Representative images in fixed gonads. (H,I) GFP fluorescence. (J,K) SNAP staining (magenta) and DAPI (cyan). n, number of germlines scored for GFP expression. Scale bar, 10 μm, in (H) applies to images. **Figure 1** and **Figure 5** results were performed in parallel, and thus results from (E,G) are the same reported in **Figure 5B,D**.

Two PGL proteins, PGL-1 and PGL-3, are the key scaffolding proteins required for P-granule assembly ^21, 22^. Genetic removal of PGL proteins causes mislocalization of P-granule proteins ^23^, aberrant expression of spermatogenic and somatic mRNAs ^15, 16, 17^, and temperature-dependent sterility ^21, 22^. PGL proteins self-assemble into granules, both *in vitro* using purified recombinant protein ^24^ and in intestinal nematode cells or mammalian cells in culture when expressed on their own ^25, 26^. Artificial PGL granules display liquid droplet behavior ^6, 11, 24^, indicating that PGL protein alone is sufficient to recapitulate the biophysical properties of P-granules in nematode cells ^10^.

PGL assembly into granules was poorly understood prior to this work though progress had been made. PGL-1 and PGL-3 are close paralogs ^22^ with the same architecture (**Figure 1B**). Many RNP granule assembly proteins rely on low complexity sequences for multivalent-multivalent, low affinity interactions ^1^. The only low complexity sequences in PGL are RGG repeats at the C-terminus, which mediate PGL RNA binding and are dispensable for granule formation ^24, 25^.

Assays for granule assembly in mammalian cells implicated the conserved N-terminal region of PGL as critical ^25^, and structural studies identified a central dimerization domain (DD) within that conserved region (**Figure 1B**) ^27^. Yet higher-ordered self-assembly demands additional PGL-PGL contacts.

In this work, we employ a tethering assay to manipulate mRNA localization in and out of granules, structural analyses to identify a new PGL dimerization domain and incisive mutational intervention to discover the role of dimerization in P-granule assembly and mRNA regulation. Our findings provide evidence that repression of mRNA expression in P-granules requires both assembly and localized regulators and hence makes a major advance in understanding the functional relationship between RNP granule assembly and function.

## Results

### Tethered PGL represses a reporter mRNA in nematode germ cells

Prior studies have suggested that P-granules regulate mRNA expression (see Introduction). To test this notion directly, we relied on a protein-mRNA tethering assay (**Figure 1C**) ^18, 19^, widely used to investigate RNA regulatory proteins ^20^. Our assay examined the expression of mRNAs to which the PGL-1 protein was tethered via λN22, a short peptide that binds with high affinity and specificity to the boxB RNA hairpin ^18^. This assay has been used successfully to identify the functions of a variety of RNA binding proteins in several organisms, including nematodes ^18, 28^. For the reporter, we inserted three boxBs into the 3’UTR of an established GFP-histone transgene that is ubiquitously expressed throughout the germline ^29^ (**Figure 1C**, Methods). To tether PGL to the GFP reporter mRNA via boxB, we generated PGL::SNAP::λN22 with sequential CRISPR gene editing in an internal, non-conserved protein region of PGL-1 (**Figures 1C** and **S1A**, see Methods). The SNAP tag ^30^ is used to visualize subcellular localization and λN22 provides tethering. PGL::SNAP::λN22 homozyotes were sterile (100%, n=94), but could be maintained and tested as a fertile heterozygote (PGL-1::SNAP::λN22/+). Given that *pgl-1* null homozygotes are fertile ^21^, the fertility defects from the addition of λN22 to PGL-1 is likely not due to defective protein function. Regardless, the logic of our tethering strategy is simple: if tethered PGL-1 localizes to granules and represses GFP expression as predicted, this assay provides a powerful entrée into fundamental questions about granule function.

To evaluate mRNA regulation, we assayed reporter GFP fluorescence in both living animals and fixed, extruded gonads; the former facilitated scoring many samples and the latter permitted scoring subcellular localization of both PGL protein via SNAP and *gfp* RNA with single molecule fluorescence *in situ* hybridization (smFISH). In controls carrying PGL-1::SNAP without λN22, GFP fluorescence was robust (**Figure 1D,F**), but GFP fluorescence was absent in animals with PGL-1::SNAP::λN22 (**Figure 1E,G**). Essentially the same result was found in fixed germlines: PGL-1::SNAP without λN22 germlines expressed GFP (100% gonads, n=35) (**Figure 1H** and **S2C,N**), but PGL-1::SNAP with λN22 had GFP was faintly detected in only a few gonads (5% gonads, n=39) (**Figure 1I** and **S2G,N**). Importantly, the PGL-1::SNAP and PGL-1::SNAP::λN22 proteins both assembled into cytoplasmic granules at the nuclear periphery (**Figure 1J,K** and **S2D,H**), similar to untagged PGL-1 and PGL-3 reported previously ^21, 22^. The SNAP signal was less for PGL-1::SNAP::λN22, perhaps because animals were heterozygous. We conclude that tethered PGL-1 localizes to perinuclear granules and represses expression of the reporter mRNA.

We next asked if the reporter RNA localized with PGL in perinuclear granules. To this end, we used smFISH to detect *gfp* RNAs and SNAP staining to detect PGL (**Figure S2A**). Control germ cells expressing PGL-1::SNAP without λN22 possessed *gfp* RNAs in both nuclear and cytoplasmic puncta (**Figure S2B-E**). We interpret nuclear puncta as nascent transcripts at active transcription sites and cytoplasmic puncta as mRNAs, based on a previous study ^31^. GFP fluorescence was robust (**Figure S2C,K**), and PGL-1::SNAP localized to perinuclear granules (**Figure S2D**), as in **Figure 1**. Germ cells expressing PGL-1::SNAP::λN22 lacked robust GFP fluorescence (**Figure S2G,K**) and PGL-1::SNAP::λN22 localized to perinuclear granules (**Figure S2H**), also as in **Figure 1**. The cytoplasmic RNA puncta were less diffuse with tethered PGL-1 than in the control and frequently colocalized with PGL-1::SNAP::λN22 in perinuclear granules (**Figure S2F-I,O**; see **Figure S3** for additional images). Imaging of the rachis, the shared cytoplasmic space in the germline, did not reveal substantial aggregates of reporter RNA as observed with mRNA in other studies ^32^. The presence of *gfp* cytoplasmic transcripts (**Figure S2F**) demonstrates that the reporter was not transcriptionally silenced, despite the lack of GFP fluorescence. In summary, reporter RNA localized with PGL-1 when tethered and PGL-1 tethering repressed reporter protein expression. These results support the idea that P-granules are sites of post-transcriptional repression.

### Identification of the PGL N-terminal Dimerization Domain

To test the significance of P-granule assembly to mRNA repression, we sought to perturb PGL-1 assembly into granules. An emerging principle is that multivalent-multivalent interactions drive granule formation (e.g. protein with at least two multimerization domains) ^1, 33^. Because PGLs are key assembly proteins for P-granules and can form granules on their own ^6, 11, 24, 25^, we reasoned that PGLs use multiple self-interactions to drive granule assembly. We previously identified one dimerization domain (DD) centrally in PGL ^27^, but DD missense mutations grossly affected protein stability. We postulated the existence of another PGL multimerization domain that might be more amenable to manipulation and again turned to structural analyses.

The region N-terminal to DD has high sequence conservation (**Figure S1B**), which implies a critical role in PGL function. Our initial efforts to express trypsin-mapped recombinant protein fragments ^27^ of this N-terminal region proved unfruitful. However, addition of amino acids disordered in the DD crystal structures ^27^ permitted robust expression sufficient for biochemical and structural characterization (**Figure S4A,B**, see Methods for more details). Henceforth, we refer to this stable protein fragment as the N-terminal dimerization domain (NtDD) (**Figure 1B**). We determined the *C. japonica* PGL-1 NtDD crystal structure to 1.5 Å (**Figure 2**, see **Table S1** for statistics, see Methods for details on crystallization and structure determination). The NtDD had a novel fold consisting of 11 alpha helices and a single N-terminal beta strand (**Figure 2B**). The asymmetric unit (ASU) was composed of four NtDD domains (**Figure 2A**), which were structurally similar (RMSD 0.219 - 0.254, chains B-D aligned to A), except for minor differences in termini and internal loops. While each ASU possessed two pairs of identical interfaces (**Figure 2A**), one of these interface pairs consisted of a network of conserved amino acid side chains making extensive salt bridges and hydrogen bonds (**Figure 3A-C** and **Figure S4C-E**). The complexity and conservation of these interactions suggested biological relevance. We next tested for dimerization *in vitro*. Recombinant PGL-3 NtDD formed a dimer on a sizing column combined with multi-angle light scattering (SEC-MALS, **Figure 3D,E**). To ask if the conserved interface in the NtDD crystal structure might be its dimerization interface, we used our structural model and *in silico* prediction ^34^ to design missense mutations predicted to disrupt the interface. These analyses yielded two distinct mutants: R123E with a single mutated residue and K126E K129E with two mutated residues. Both NtDD interface mutants formed monomers rather than dimers in solution (**Figure 3D,E**). We conclude that the dimers observed in the crystal structure represent the NtDD dimer detected in solution. Therefore, PGL proteins possess two dimerization domains. We renamed the original DD to Central DD (CDD) for clarity (**Figure 1B**).

**Figure 2.**
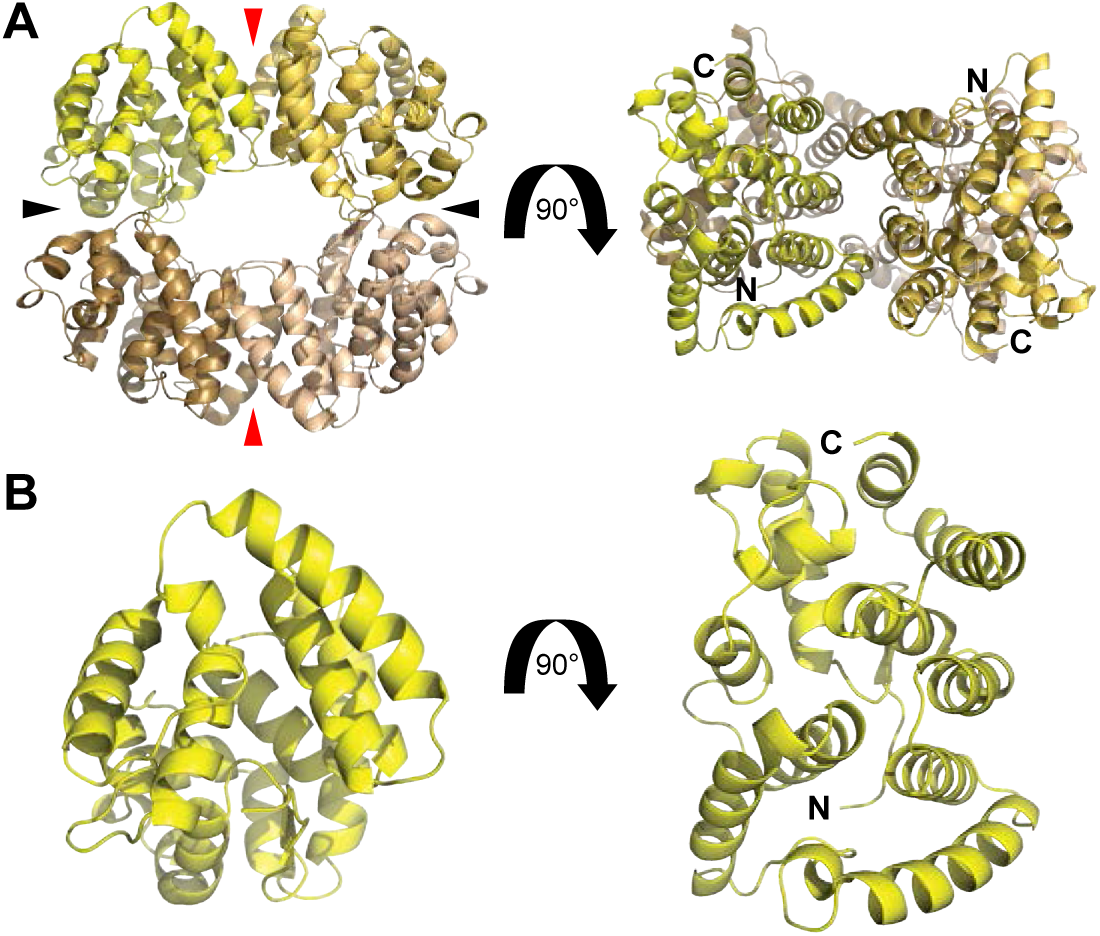
Crystal structure of PGL NtDD (A) Crystal structure of *C. japonica* PGL-1 NtDD to 1.5 Å. See **Table S1** for crystal structure data and model statistics. NtDD has four copies per asymmetric unit (ASU). Copies in yellow, gold, tan, and brown. Arrows indicate two pairs of subunit interfaces in the ASU. Red arrows highlight the interface relying on conserved amino acids (see text). (B) Enlarged image of a single NtDD.

**Figure 3.**
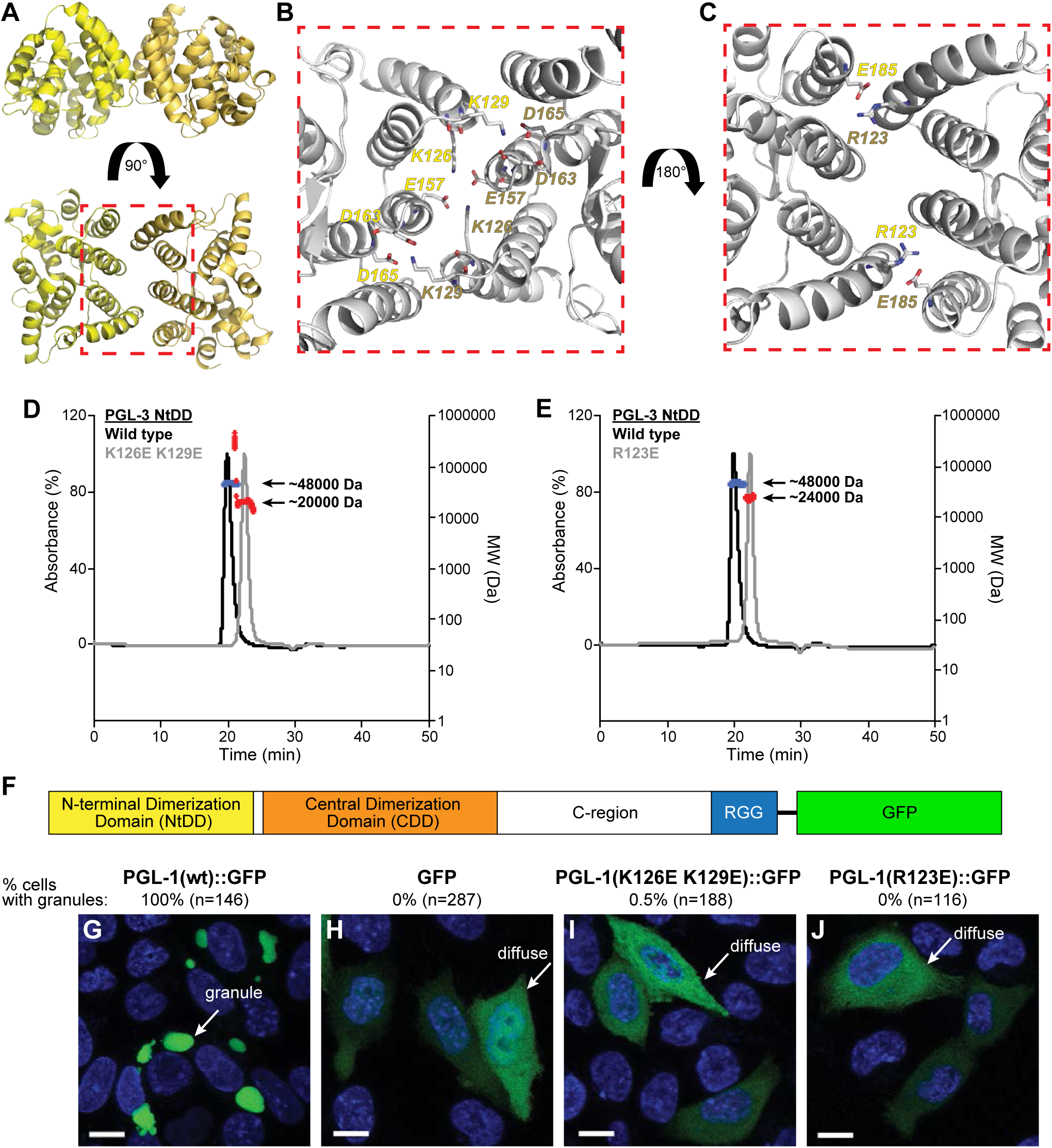
NTD dimerization and its role in PGL self-assembly (A) Structural model of the NtDD dimer. (B-C) Enlargement of dimer interface (red box in A). PGL-1 amino acids (B) K126 and K129, and (C) R123 interact with apposing subunit side chains. Residue labels in yellow or gold to indicate their representative subunits. (D,E) Size exclusion chromatography and multi-angle light scattering (SEC-MALS) of recombinant PGL-3 (D,E) NtDD wild type, (D) K126E K129E and (E) R123E proteins. A280 UV absorbance (left y axis) was normalized to the maximum value. Molecular weight (MW, right y axis) for MALS in daltons (Da). Wild-type protein (blue) measured the approximate size of a dimer, while both mutant proteins (red) measured approximately as monomers. (F) Diagram of *C. elegans* PGL-1 C-terminally tagged with GFP. (G-J) Representative images of (G) GFP-tagged PGL-1, (H) GFP alone, and GFP-tagged PGL-1 (I) K126E K129E and (J) R123E mutants expressed in Chinese Hamster Ovary (CHO) cells. Cell cultures were imaged live, and GFP-positive cells counted for the presence or absence of granules. Images show the majority result (percentages noted above image). Scale bar, 10 µm.

### PGL NtDD dimerization is critical for granule formation

To assess the role of NtDD dimerization in PGL granule self-assembly, we used an assay in mammalian cells where expression of wild-type PGL-1 tagged with GST was sufficient for assembly into granules ^25^. Similar to that report, wild-type PGL-1 tagged with GFP also formed large cytoplasmic granules when expressed in mammalian cells (**Figure 3F,G**), while GFP alone was diffuse (**Figure 3H**). However, PGL-1::GFP mutated to either K126E K129E or R123E no longer self-assembled into granules (**Figure 3I,J**). We conclude that NtDD dimerization is essential for granule formation in mammalian cells.

To assess the role of NtDD dimerization in nematode germ cells, we used CRISPR to introduce the dimerization defective mutations into SNAP-tagged PGL-1 (**Figure 4A**, **Figure S1A**, Methods). We first asked about effects on fertility (**Figure 4B**). Most wild-type PGL-1 (no SNAP) and PGL-1::SNAP animals were fertile at 20°C and 25°C (**Figure 4B**). In contrast, most *pgl-1* null mutants were fertile at 20° but few were fertile at 25°C (**Figure 4B**), as reported previously ^21, 22^. Fertility of the NtDD dimerization mutants, K126E K129E and R123E, by contrast, was sharply reduced at both 20° and 25°; many mutant worms were sterile at 20°C and most were sterile at 25°C (**Figure 4B**). Fertility was therefore impacted more severely in dimerization mutants than *pgl-1* null mutants (**Figure 4B**), suggesting that a PGL mutant protein incapable of NtDD dimerization functions as a dominant-negative. The NtDD dimerization mutants had smaller than normal germlines and many lacked oocytes (**Figure S5B-D**), which are defects typical of *pgl-1* and *pgl-1 pgl-3* null mutants ^21, 22^. We conclude that PGL-1 NtDD dimerization is critical for fertility.

**Figure 4.**
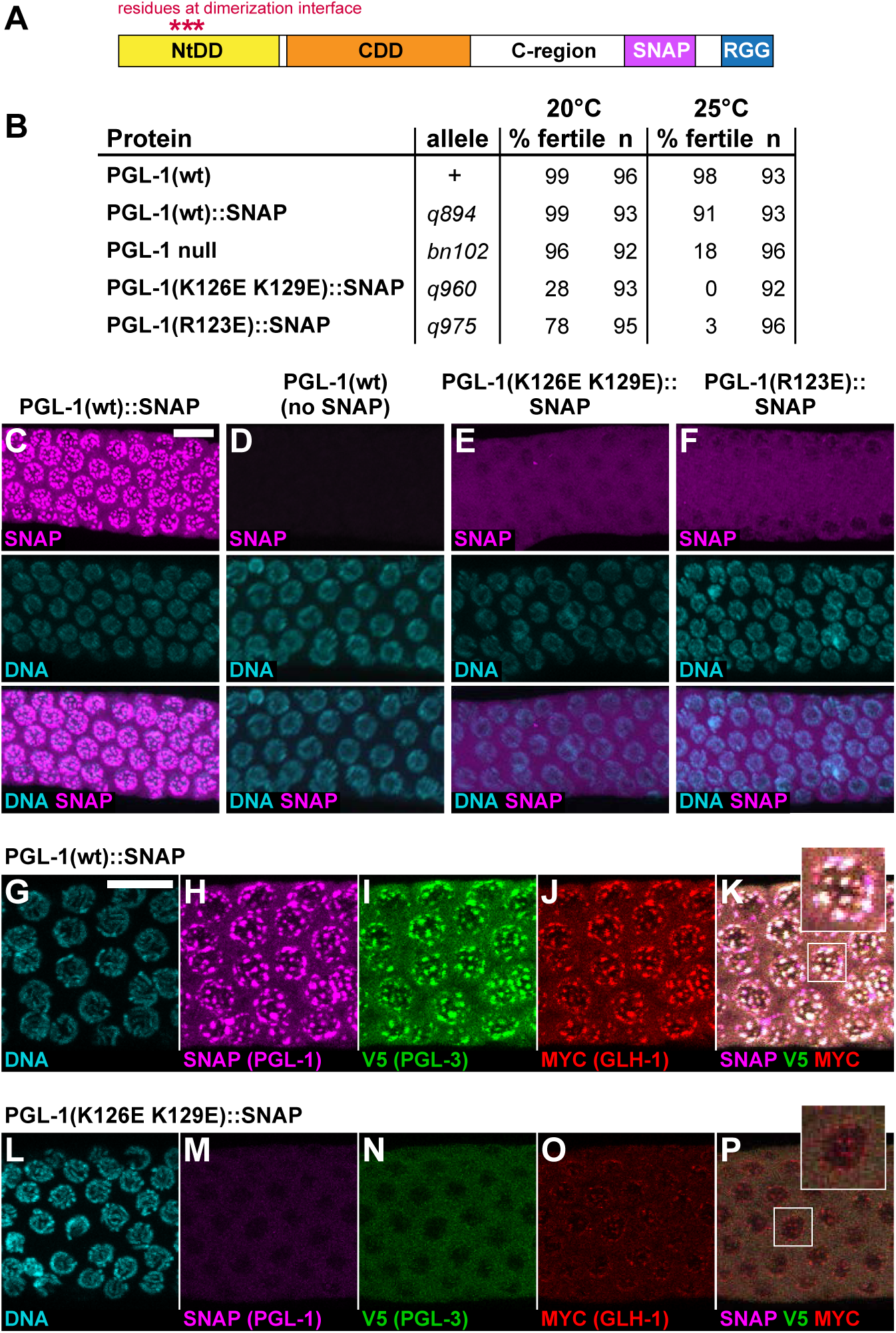
NtDD dimerization is critical for fertility and P-granule formation in nematodes (A) Sites of SNAP tag insertion and missense mutations in *C. elegans* PGL-1. (B) Fertility of SNAP-tagged PGL-1 animals. Percentages were obtained after scoring individuals for production of larval progeny after 5 days at either 20°C or 25°C. (C-P) Extruded adult germlines, fixed, stained and imaged in same region of meiotic pachytene (see **Figure S5A**). (C-F) Representative images of SNAP staining to visualize PGL-1 expression and granule formation. All images are partial z-stacks to maximize visualization of P-granules. Images were taken from germlines containing embryos; similar images were obtained from germlines lacking embryos (**Figure S5E,F**). (C) PGL-1::SNAP localizes to granules around nuclei (n=49). (D) Control, wild-type animal lacking SNAP tag shows virtually no background staining (n=20). (E) PGL-1(K126E K129E)::SNAP is diffuse (n=38). (F) PGL-1(R123E)::SNAP is diffuse (n=24). (G-P) Representative images showing localization of three P-granule components in germ cells expressing either (G-K) PGL-1::SNAP (n=20) or (L-P) PGL-1(K126E K129E)::SNAP (n=14). (G,L) DNA (DAPI); (H,M) SNAP (PGL-1::SNAP or mutant); (I,N) V5 (PGL-3); (J,O) MYC (GLH-1); (K,P) Merge. Scale bar, 10 µm for all images, except 2.5-fold enlargements of nuclei in boxes placed outside main images.

To investigate the role of PGL NtDD dimerization in granule assembly, we compared the subcellular localization of wild-type PGL-1::SNAP to NtDD dimerization-defective PGL-1::SNAP mutant proteins (**Figure 4C-F**). Wild-type PGL-1::SNAP assembled into cytoplasmic granules at the nuclear periphery (**Figure 4C**), but the K126E K129E and R123E mutant proteins were largely diffuse in both fertile (**Figure 4E,F**) and sterile (**Figure S5E,F**) germlines. Protein expression levels may affect the propensity of PGL-1 to form granules but were difficult to compare due to variability in germline size. Mutant PGL-1 fluorescent intensity was above background (**Figure S6A-E**), albeit less than wild-type PGL-1. Immunoblots also revealed only a modest difference in protein expression between PGL-1 wild-type and mutant protein (**Figure S6F**). Despite these differences, the mutant PGLs were capable of forming small perinuclear granules in a variable number of germ cells (**Figure 4E,F** and **Figure S5E,F**). In addition, for each mutant, we found a single germline (1 of 59 for K126E K129E; 1 of 54 for R123E) with PGL-1 perinuclear granules in all germ cells (**Figure S5G,H**). Therefore, both PGL-1 mutant proteins are capable of incorporating into granules, but do so much more inefficiently than their wild-type counterparts (**Figure 5C**).

**Figure 5.**
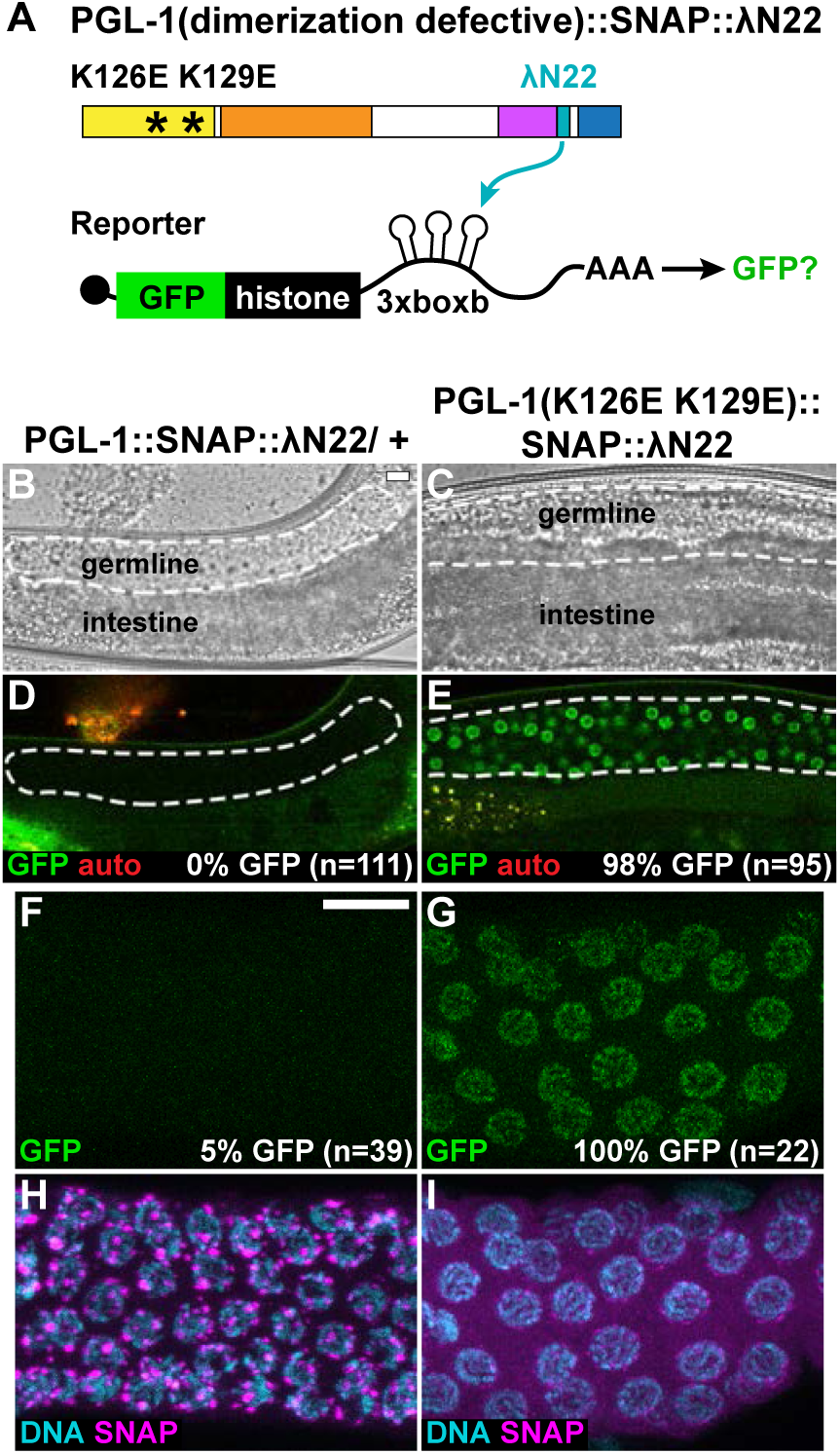
PGL assembly is required for tethered mRNA reporter repression (A) Protein-mRNA tethering assay and PGL assembly. To test the necessity of granule formation for mRNA repression, NtDD assembly mutations were added to PGL-1::SNAP::λN22 and germlines observed for GFP reporter expression. (B-E) GFP reporter expression in germ cells of live animals. (B,C) brightfield image. (D,E) GFP fluorescence (green); auto fluorescence (red). n, number of animals scored for GFP expression. Scale bar, 10 μm, in B applies to B-E images. (F-I) Representative images of PGL granule formation, seen by SNAP staining (magenta) and GFP fluorescence (green) in fixed gonads. n, number of germlines scored for GFP expression. “%”, germlines with detectable GFP. Scale bar, 10 μm, in F applies to F-I images. **Figure 1** and **Figure 5** results were performed in parallel, and thus results from B,D are the same reported in **Figure 1E,G**.

We next asked why the fertility defects of PGL-1 NtDD dimerization mutants were more severe than a *pgl-1* null mutant. The likely explanation was interference with assembly of other P-granule components into granules. Normally, PGL-1 interacts with PGL-3 ^22^, and both PGL-1 and PGL-3 rely on GLH-1 or GLH-4 Vasa helicases to localize to the nuclear periphery in adult germ cells ^26, 35^. In contrast, GLH proteins can assemble at the nuclear pore independently of PGLs ^22^. We postulated that PGL-1 assembly mutants might interfere with PGL-3 assembly into granules but not affect GLH-1. To test this idea, we epitope-tagged endogenous *pgl-3* and *glh-1* (see Methods) and compared localization of PGL-3::V5 and GLH-1::Myc in germ cells also expressing either wild-type PGL-1::SNAP or the dimerization defective mutant PGL-1(K126E K129E)::SNAP. In the presence of wild-type PGL-1::SNAP, all three proteins, PGL-1, PGL-3 and GLH-1, co-localized to granules at the nuclear periphery (**Figure 4G-K**), as previously observed for untagged proteins ^21, 22, 36^. By contrast, a dimerization-defective mutant protein, PGL-1(K126E K129E)::SNAP, was diffuse rather than granular, and wild-type PGL-3 became similarly diffuse (**Figure 4L-N**). GLH-1, however, was still capable of localizing to granules at the nuclear periphery (**Figure 4O-P**), similar to previous reports ^22^. Although these GLH-1 granules appeared smaller than normal, their formation was seen around virtually all germline nuclei (**Figure 4P**). We conclude that NtDD dimerization of PGL-1 is crucial for assembly of both PGL-1 and PGL-3 into granules and that this likely explains the severe fertility defects of PGL-1 dimerization mutants.

### PGL granule assembly is required for PGL-mediated mRNA repression

The identification of assembly-defective PGL-1 proteins coupled with our tethering assay (**Figure 5A**) allowed us to test the relationship between granule assembly and mRNA repression. We introduced the K126E K129E mutation into PGL-1::SNAP::λN22 and tested for reporter expression. PGL-1(K126E K129E)::SNAP::λN22 homozygotes were fertile (21%, n=96) to an extent comparable to PGL-1(K126E K129E)::SNAP without λN22 (**Figure 4B**). While control PGL-1::SNAP::λN22 repressed the reporter (**Figure 5D,F**), assembly-defective PGL-1 was not repressive and the vast majority of germ cells expressed GFP (**Figure 5E,G and S2K,N**). The PGL-1(K126E K129E)::SNAP::λN22 mutant protein was diffuse and non-granular compared to wild-type PGL-1::SNAP (**Figure 5H,I**), similar to PGL-1(K126E K129E)::SNAP without λN22 (**Figure 4E**). By smFISH, *gfp* RNA signal in PGL-1(K126E K129E)::SNAP::λN22 mutant germlines was observed robustly throughout the cytoplasm (**Figure S2J-M**). Formally, the PGL-1 interface residues might affect granule assembly and mRNA repression independently but the simplest explanation is that PGL-1 must assemble into granules to repress mRNA.

### PGL-mediated mRNA repression relies on WAGO-1, a cytoplasmic Argonaute

PGL tethering provides a simple assay for identification of additional factors needed for P-granule mRNA repression (**Figure S7A**). We depleted candidate P-granule-associated RNA regulators with RNAi and sought GFP reporter de-repression in PGL-1::SNAP::λN22 worms. In this candidate screen, knockdown of the cytoplasmic Argonaute WAGO-1 had a dramatic effect (**Figure S7B**). RNAi against other candidates, by contrast, had either no or a minor effect on GFP repression (**Figure S7B**). The RNAi screen highlighted WAGO-1 as a key factor in PGL-mediated mRNA repression, a finding consistent with previous studies showing that WAGO-1 localizes to P-granules and regulates gene expression ^37, 38^.

To further investigate *wago-1*, we first inserted an epitope-tag at the endogenous locus (**Figure S7C**, Methods). WAGO-1::3xV5 co-localized with PGL-1(wt)::SNAP in perinuclear granules (**Figure 6A**), consistent with a previous report that WAGO-1 resides in P-granules ^37^. We next asked if WAGO-1 association with perinuclear granules was dependent on PGL assembly. In animals expressing assembly-defective PGL-1(R123E)::SNAP, which fails to form granules efficiently (**Figure 4E,F**), WAGO-1 was diffuse in about half the germlines (**Figure 6B**), but granular in the other half (**Figure 6C**). Therefore, WAGO-1 can assemble into P-granules independently of PGL-1. To investigate if PGL assembly is independent of WAGO-1, we generated an internal deletion that creates a frameshift and fails to express WAGO-1 protein (**Figure 6D**, **Figure S7C**, Methods). PGL-1::SNAP localized to perinuclear granules in the absence of WAGO-1 (**Figure 6D**). In sum, PGL-1 and WAGO-1 can assemble into granules independently of each other.

**Figure 6.**
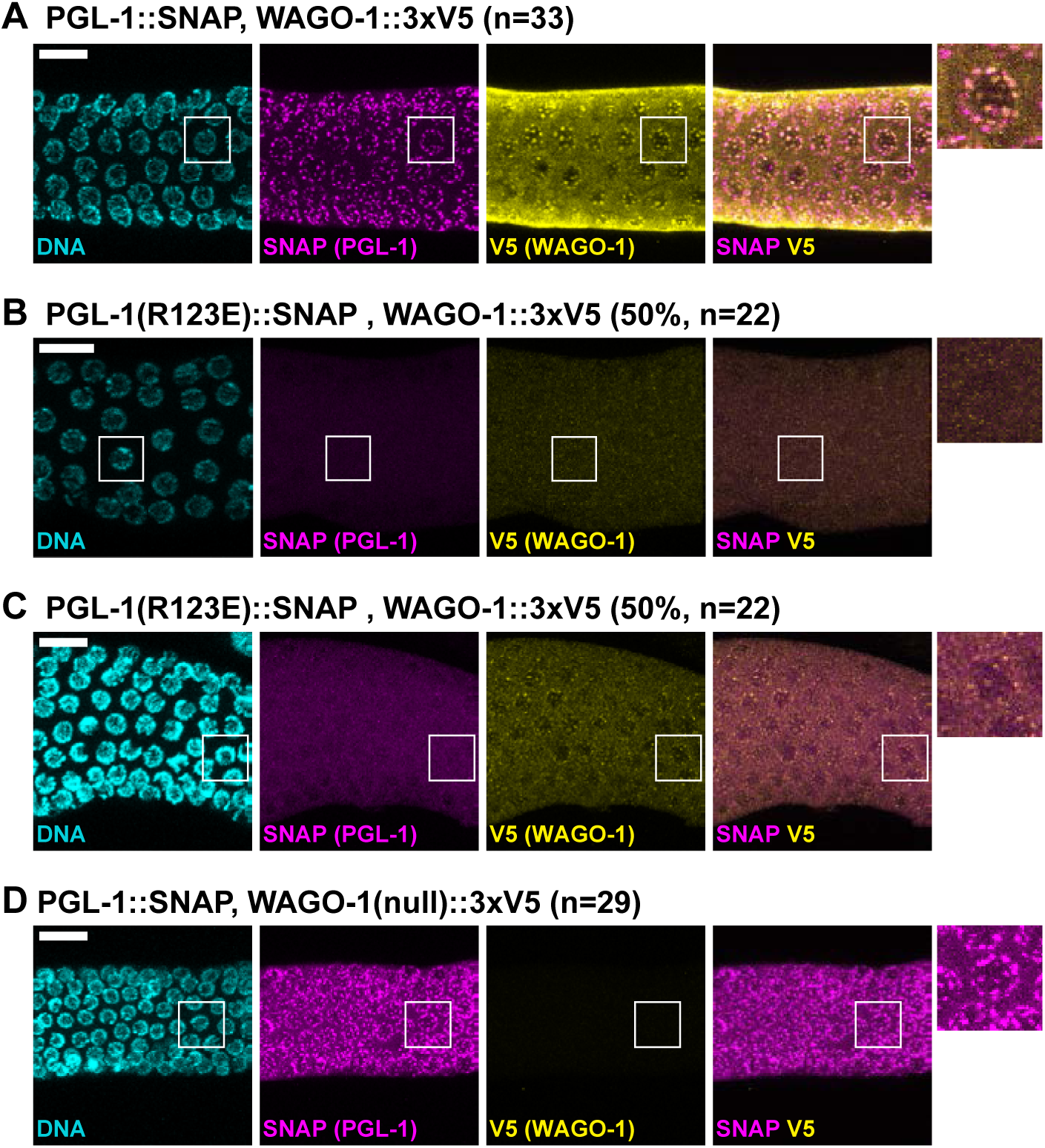
PGL-1 and WAGO-1 assemble independently in P-granules (A-D) Representative images showing localization of PGL-1 and WAGO-1 in germ cells expressing (A) PGL-1::SNAP, WAGO-1::3xV5 (n=33). (B) PGL-1(R123E)::SNAP, WAGO-1::3xV5 without WAGO-1 puncta (11 of 22 germlines). (C) PGL-1(R123E)::SNAP, WAGO-1::3xV5 with WAGO-1 puncta (11 of 22 germlines). (D) PGL-1::SNAP, WAGO-1(null)::3xV5 (n=29). DNA (DAPI); SNAP (PGL-1::SNAP or mutant); V5 (WAGO-1::3xV5 wild-type or null). Scale bar, 10 µm for all images, except 2.5-fold enlargements placed outside main images.

We finally explored the role of WAGO-1 in mRNA repression within granules (**Figure 7A**). To this end, we tested PGL-1::SNAP::λN22 for its ability to repress reporter RNA in the presence or absence of WAGO-1. We again conducted our assays in living animals to ensure a large sample size (**Figure 7B-E**) and fixed extruded gonads to visualize PGL in addition to GFP fluorescence (**Figure 7F-I**). The reporter was repressed with wild-type WAGO-1 (**Figure 7D,F**), as expected, but de-repressed in the *wago-1* null mutant (**Figure 7E,G**), consistent with *wago-1* RNAi (**Figure S7B**). The de-repression was seen in ∼70% of germlines when assayed in living animals (**Figure 7E**) and in ∼90% of fixed gonads (**Figure 7G**). This more penetrant de-repression in fixed gonads might reflect higher sensitivity or lower sample size. Loss of *wago-1* had no observable effect on fluorescent reporter expression (**Figure S8**), arguing against varying reporter mRNA levels causing the increase in fluorescence. Regardless, the percentages demonstrate strong but incomplete de-repression, a result consistent with WAGO-1 functioning redundantly with other Argonautes ^37, 38, 39^. As expected, PGL-1::SNAP::λN22 assembles into granules, with or without WAGO-1 (**Figure 7H,I**). Moreover, by smFISH, the *gfp* reporter mRNA colocalizes with PGL-1 both in animals with wild-type WAGO-1 (**Figure S9A-D,I**) and in the *wago-1* null mutant (**Figures S9E-I** and **S10**). We conclude that WAGO-1 is a major regulator of mRNA repression in P-granules. These results also provide direct evidence that granule formation alone is not sufficient for mRNA regulation (see **Figure 8**, Discussion).

**Figure 7.**
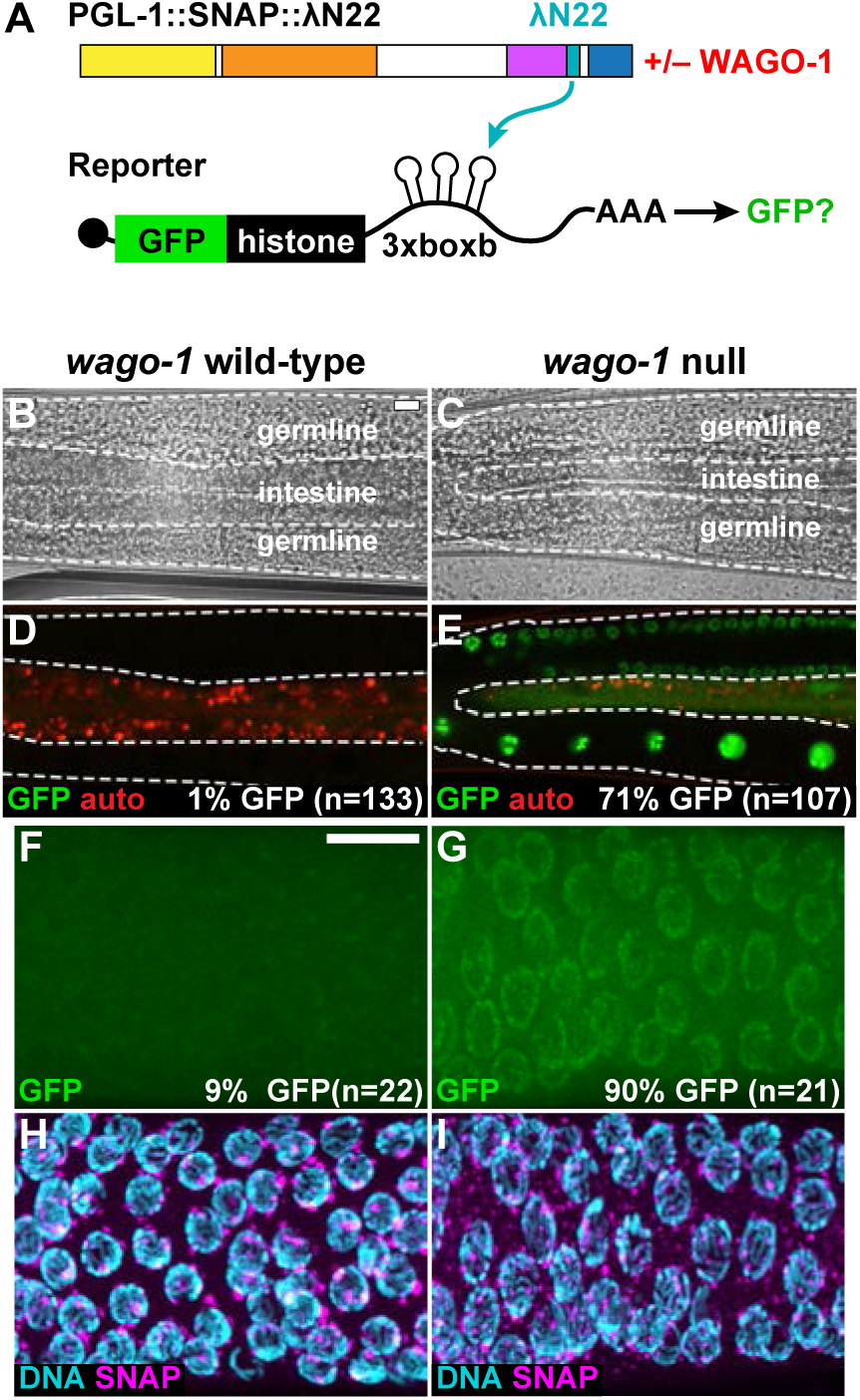
Argonaute WAGO-1 is required for PGL-tethered mRNA reporter repression (A) Protein-mRNA tethering assay and WAGO-1. To test the necessity of WAGO-1 for mRNA repression, PGL-1::SNAP::λN22 germlines were analyzed for GFP reporter expression in the presence or absence of WAGO-1. (B-E) GFP reporter expression in germ cells of live animals with (B,D) wild-type *wago-1* or (C,E) *wago-1* null. (B,C) Brightfield image. (D,E) GFP fluorescence (green); auto fluorescence (red). n, number of animals scored for GFP expression. Scale bar, 10 μm, in B applies to B-E images. (F-I) Representative images of fixed germlines with (F,H) wild-type *wago-1* or (G,I) *wago-1* null. (F,G) GFP fluorescence. (H,I) DNA (DAPI) and PGL granule formation seen by SNAP staining. n, number of germlines scored for GFP expression. Scale bar, 10 μm, in F applies to images F-I.

**Figure 8.**
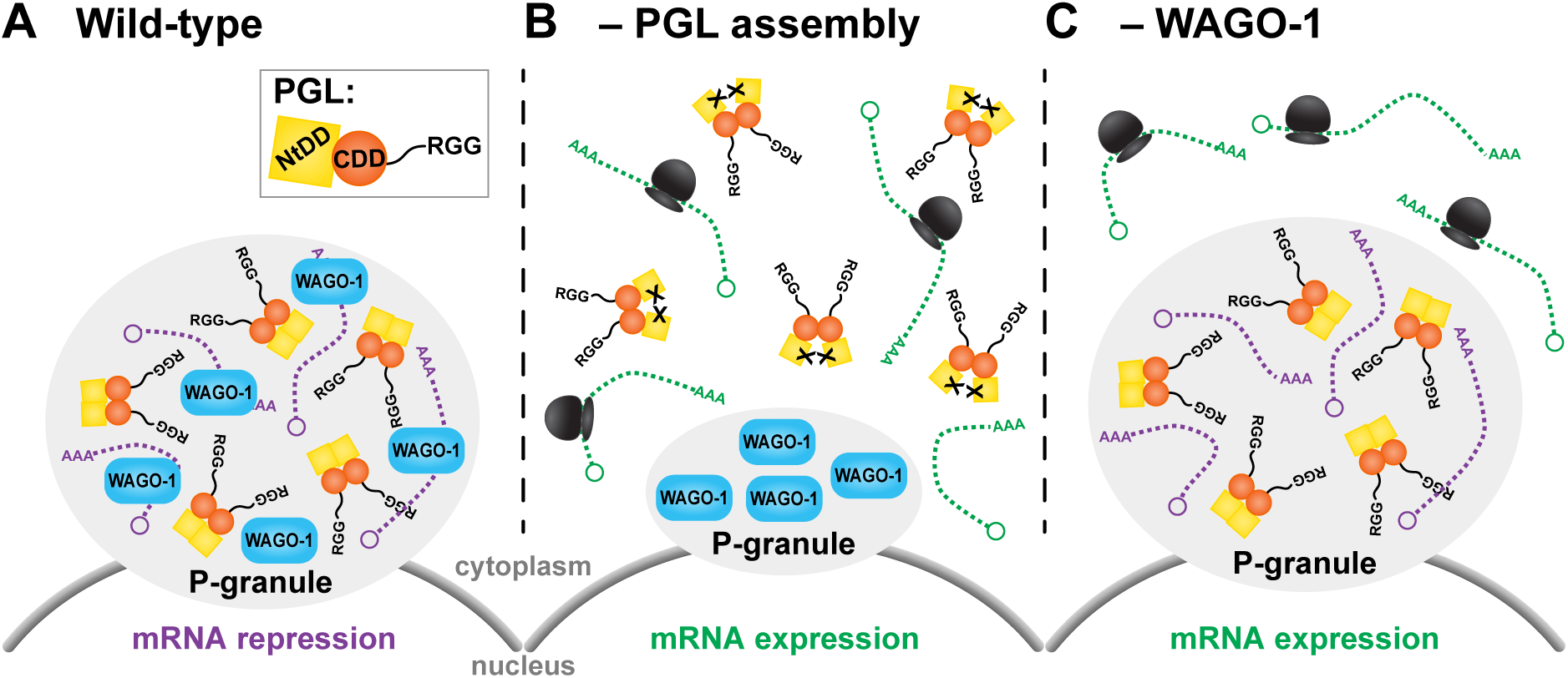
Model of P-granule assembly and mRNA repression (A) P-granules assemble at the nuclear pore with assembly-competent PGL protein. PGL is shown as a dimer for simplicity but multivalent PGLs likely form an oligomeric protein-RNA network. WAGO-1 binds to RNA, as expected for an Argonaute, and WAGO-associated RNAs are repressed through mRNA turnover or translational repression (purple). (B) When PGL NtDD cannot dimerize, PGL fails to assemble into P-granules at the nuclear periphery. WAGO-1 assembles independently into P-granules. PGL-associated, non-granular mRNAs are available for translation (ribosomes, black). (C) In the absence of cytoplasmic Argonaute WAGO-1, PGL proteins assemble into P-granules normally with its associated mRNA, but loss of WAGO-1 perturbs repression of P-granule-localized transcripts. PGL’s liquid droplet properties permit diffusion of some transcripts into the cytoplasm for translation. RNAs in the granule are not translated, either because of repressors or the lack of ribosomes. See text for further Discussion.

## Discussion

This work investigates the functional relationship between assembly of a paradigmatic liquid droplet, the *C. elegans* P-granule, and the activities of its regulatory components. Our analyses make three key advances. First, we discover that PGL-1 dimerization of the N-terminal domain is required for its assembly into P-granules. Second, we demonstrate that PGL-mediated mRNA repression relies on PGL granule assembly. Third, we find that mRNA repression by P-granules employs the activity of at least one P-granule constituent, the Argonaute WAGO-1. Together, these advances support a model that PGL assembly into granules is necessary for its biological function but not sufficient to repress the expression of localized mRNAs (**Figure 8**). Below we discuss these advances and their implications for RNP granules more generally.

### Dimerization drives PGL protein assembly into granules

An emerging principle of RNP granule assembly is that multivalent macromolecular interactions drive granule formation ^1, 33^. This work extends that principle to nematode P-granules and their primary assembly proteins, the PGLs (see Introduction). A previous study reported that PGL proteins possess a dimerization domain (DD) in their central region ^27^, and here we report the discovery of a second PGL DD in the N-terminal region (NtDD). Thus, PGLs possess two protein folds that confer multivalency. Based on insights from the NtDD structure, we designed two distinct PGL-1 mutant proteins that are unable to dimerize *in vitro* and are severely compromised *in vivo* for assembly into P-granules and fertility. These results provide evidence that PGL dimerization is crucial for P-granule assembly and for P-granule biological function. Although attempts to disrupt dimerization via the central DD led to protein instability and hence could not test its biological significance ^27^, we include both N-terminal and central DDs in our model for how PGLs mediate assembly into P-granules (**Figure 8A**). A critical future direction is to investigate how each dimerization domain contributes to higher order and likely oligomeric assembly.

Self-assembling, multivalent proteins have been identified for several RNP granules. Examples include TDP-43 in nuclear granules ^40^, Oskar and Vasa for *Drosophila* polar granules ^41, 42, 43^; EDC3 and LSM4 for P-bodies ^44^; and MEG-3 and MEG-4 for embryonic P-granules ^6, 45, 46^. These various assembly proteins rely on a combination of multimerization domains and low complexity, intrinsically disordered sequences to facilitate granule formation ^6, 44, 45, 47, 48, 49^. A leading hypothesis of liquid droplet assembly invokes reliance on multiple, weak interactions among RNP constituents ^1, 33^. Our work highlights the idea that interactions between structured regions are a driving force *in vivo* for assembly of liquid droplet granules as well as their biological function (see below). Recombinant PGL proteins make liquid droplets on their own *in vitro* ^6, 11, 24^, suggesting that PGLs possess regions responsible for low affinity contacts in the full-length protein. We suggest that PGL uses both dimerization domains as well as additional low affinity contacts to facilitate liquid droplet formation. Alternatively, the DDs may be subject to post-translational modifications that modulate their affinity *in vivo*. PGL did not require its C-terminal RGG repeats to form granules in mammalian culture ^25^, but RGG repeats in other assembly proteins enhance granule formation ^49^. The PGL RGG repeats may instead be needed to trigger robust granule assembly with RNA ^24^ or impart liquid droplet properties associated with PGL in nematodes ^10^. Regardless, the discovery that PGL proteins are multivalent with two dimerization domains is an essential advance for understanding the assembly of this paradigmatic liquid droplet.

### PGL-mediated mRNA repression relies on recruitment into P-granules

Our study employs a protein-RNA tethering assay to provide direct evidence that mRNA expression is repressed when the mRNAs are recruited to P-granules. Three lines of indirect evidence had previously led to this idea: P-granules contain repressed mRNAs ^12^; spermatogenic and somatic transcripts are aberrantly expressed in the absence of P-granules ^15, 16, 17^; and a variety of RNA regulatory factors co-localize in P-granules, including inhibitory RNA binding proteins, like the Pumilio homolog FBF-2 ^32^, and RNA regulatory enzymes, like the Argonaut/Piwi PRG-1 ^50^ and the deadenylase PARN-1 ^51^. Despite these clues, the significance of P-granule assembly for biological function remained enigmatic. Our discovery of key amino acids required for PGL assembly into granules (see above) allowed us to compare RNA regulation when coupled to assembly-competent or assembly-defective PGL. We found that a reporter mRNA tethered to assembly-competent PGL protein localized with PGL to P-granules and was repressed, but a reporter tethered to an assembly-defective PGL did not localize to granules and was expressed. Together, these results suggest that localizing mRNAs into P-granules leads to their repression (**Figure 8A,B**).

### Argonaute WAGO-1 promotes repression of mRNA expression in P-granules

Expression of mRNAs localized within an RNP granule might be repressed by either of two broad mechanisms. Granule localization might recruit RNAs to form interactions with negative-acting factors (e.g. RNA turnover machinery) or it might sequester RNAs away from positive-acting factors (e.g. translational machinery). Our results suggest that localization with assembled PGL and key repressors is the primary mechanism (**Figure 8**). Previous work identified an *in vitro* RNase activity for PGL DD ^27^. However, PGL-tethering of RNA, and subsequent recruitment into granules, is not sufficient for repression on its own but also relies on a P-granule-localized Argonaute, WAGO-1 (**Figure 8C**). Previous studies reported that WAGO-1 is P-granule-associated and represses transcripts that have been primed by the piRNA pathway as part of the secondary RNAi response ^38^. Our tethering results broaden the role of WAGO-1 to repress any RNA associated with the P-granule scaffold. Previous studies also reported that WAGO-1 is functionally redundant with other Argonautes ^37, 38, 39^. We suspect this redundancy may explain why some, not all, *wago-1* null mutant germlines repressed PGL-tethered reporter mRNAs. The specific mechanism for mRNA repression, mRNA turnover or translational repression, is still unclear, although we favor RNA turnover driven by PGL’s RNase activity and the Argonaute. Additional work must be done to understand the relative importance and individual roles of these Argonautes and PGL in mRNA repression.

Other cytoplasmic RNP granules have been proposed to repress mRNAs. Granule formation of a yeast amyloid-like RNA binding protein correlates with translational inhibition of transcripts critical for gametogenesis ^52^, and P-bodies contain mRNAs that are translationally repressed in cells ^53^. Our work extends this idea further by showing the dependence of other enzymatic factors in liquid droplet granules for mRNA repression. Our model proposes that loss of PGL dimerization prevents P-granule assembly and hence compromises the coalescence of RNAs with WAGO-1 and other P-granule-associated repressors (**Figure 8B**), while loss of WAGO-1 relieves granule-recruited mRNAs from repression (**Figure 8C**). Therefore, the scaffolding function of PGL is important but not sufficient for P-granule function. We also propose that, without WAGO-1, mRNAs that are normally destined for P-granule association can exist in either of two states. Those remaining in the granule are not expressed due either to the presence of other RNA repressors or the absence of ribosomes, but those diffusing out of the liquid droplet granule reach the translational machinery for expression (**Figure 8C**). By extension, we propose that other RNP granules with liquid droplet properties may require both scaffolding proteins and active enzymatic factors to regulate their mRNAs.

This work adds to the emerging theme that granules play a general role in regulating the function of their components. For example, stress granules sequester the mTORC1 protein complex to block activation of mTOR signaling ^54^, mammalian cells can trap hormones and melanin in amyloid-like aggregates to prevent active signaling ^55, 56^, and liquid droplet granule formation of cGAS with cytosolic DNA triggers its enzymatic activity *in vitro* and cell culture ^57^. The functional relationship between PGL and WAGO-1 is unknown. PGL may simply recruit or retain mRNAs in P-granules to be regulated by WAGO-1. A more enticing model is that PGL forms a higher ordered, RNP complex with mRNA that enhances WAGO-1’s biochemical activity. Further studies that pair insights into mechanisms of granule assembly with direct *in vivo* assays of regulation will be pivotal moving forward to decipher the mechanistic function of other RNP granules in their biological context.

## Materials and Methods

### Protein expression and purification

We previously used *C. elegans* PGL-3 recombinant protein and limited proteolysis to identify a central dimerization domain (CDD) ^27^. While we could express CDD efficiently we could not express recombinant protein that was N-terminal to the cleavage site (PGL-3 amino acid residues 205-206). We tried moving the six-histidine purification tag to the N- and C-termini, shortened the protein regions used for expression, and tried several different orthologs with little success. The insight came after aligning protein sequences of several Caenorhabditid sp. and studying the CDD domain boundary (**Figure S1B**). Protease cleavage occurred in a conserved portion of the N-terminal region and this region was disordered in our DD crystal structures. After inclusion of this region (PGL-3 amino acid residues 205-212), we could express and purify recombinant N-terminal protein from *C. elegans* PGL-1 and its orthologs. We refer to this region as the N-terminal domain (NtDD).

This study used primarily *C. elegans* PGL-3 and *C. japonica* PGL-1 recombinant NtDD proteins. The *C. elegans* PGL-3 coding region was PCR amplified from cDNA. A codon-optimized (*E. coli*) version of *C. japonica* PGL-1 NtDD was ordered as a gBlock (IDT, Coralville, IA). We included a six-histidine tag at the C-terminus that was removed later with carboxypeptidase A ^58^. Constructs were cloned into a pET21a vector (MilliporeSigma, Burlington, MA) using Gibson Assembly cloning ^59^, and plasmids transformed into Rosetta2 cells (MilliporeSigma, Burlington, MA). Cultures were grown at 37°C with shaking (225 rpm) until ∼0.8 OD, cooled for 30-60 minutes, and induced with a final concentration of 0.1 mM IPTG. Cultures were then grown at 16°C with shaking (160 rpm) for 16-18 hours, collected, and bacterial pellets frozen until use. Selenomethionine-incorporated *C. japonica* protein was expressed in SelenoMethionine Medium Complete (Molecular Dimensions, Suffolk, UK), and grown, induced, and collected in a similar manner.

Bacterial pellets were defrosted on ice and reconstituted in lysis buffer (20 mM sodium phosphate pH 7.4, 300 mM NaCl, 10 mM imidazole, 5 mM beta-mercaptoethanol (BME)) with protease inhibitors (cOmplete™ EDTA-free, Roche, Indianapolis, IN). Lysozyme (Sigma-Aldrich, St. Louis, MO) was added at 50 µg/ml and incubated on ice for 20 minutes prior to lysis in a french press. Samples were spun at low (3220 × g, 4°C, 20 minutes) and high speed (10,000 × g, 20°C, 10 minutes), then incubated with 1.5 ml NiNTA beads (Thermo Fisher Scientific, Waltham, MA) for 1 hour at 4°C with rotation. Sample supernatant was separated by gravity flow, washed twice with lysis buffer, and eluted using lysis buffer with increasing imidazole concentrations (20, 40, 60, 80, 100, 250 mM). Eluted samples were checked for protein via Bradford assay (Bio-Rad, Hercules, CA), and dialyzed overnight in HN buffer (20 mM HEPES pH 7.4, 100 mM NaCl). The dialyzed samples were concentrated with a Centriprep 10K concentrator (Millipore), calcium added to 1 mM CaCl_2_, and the histidine tag removed with carboxypeptidase A bound to agarose (Sigma, St. Louis, MO) at a ratio of 10 protein:1 enzyme (w/w). Samples were incubated at room temperature (∼20°C) for 45-90 minutes with rotation prior to supernatant elution by centrifugation in microflow columns (Thermo Fisher Scientific, Waltham, MA). Samples were run on a S200 sizing column (GE Healthcare, Chicago, IL) in HNT buffer (20 mM HEPES pH 7.4, 100 mM NaCl, 0.5 mM TCEP pH 7.4). Fractions containing recombinant protein were collected, concentrated in an Amicon 10K concentrator (MilliporeSigma, Burlington, MA), and protein concentration estimated by A280. Samples were frozen in liquid nitrogen or used immediately.

### Crystallization and structure determination

*C. elegans* PGL-1, *C. elegans* PGL-3, and *C. japonica* PGL-1 NTD recombinant protein were screened in crystallization conditions using 400 nl hanging and sitting drop 96-well trays set up with the Mosquito (TTP Labtech, Cambridge, MA) in 20°C. Several conditions produced labile crystal plates. Data was collected to 4 Å from *C. elegans* PGL-1 crystal plates, determined to have a very large unit cell (86 Å × 86 Å × 460 Å) and P6 point group, and eventually determined to have perfect merohedral twinning. *C. japonica* PGL-1 also crystallized as large (60-150 Å) rhomboid crystals in 40-45% PEG 400 at low (Na Citrate pH 5.5-6.0) and physiologic pH (imidazole pH 7.5-8.0). Crystals grown in citrate or imidazole both diffracted well, but we used imidazole (100 mM imidazole pH 7.5, 45% PEG 400, 1 mM TCEP pH 7.4) due to its higher reproducibility for large crystals and its modestly better resolution. The crystals did not require additional cryo-protection due to the high PEG 400. We eventually collected a full data set to 1.5 Å in space group C2.

PGL-1 NtDD was a novel domain. Novelty and translational pseudosymmetry precluded us from using any model for molecular replacement. Trial heavy atom soaks also proved unfruitful, and the *C. japonica* PGL-1 NtDD has just two methonines past the start codon, making selenomethionine phasing challenging. To boost anomalous signal, we mutated two non-conserved isoleucines to methionines (I63M, I212M). This methionine mutant provided phases to 3.6 Å by single anomalous dispersion (SAD) that we used to build a 1.6 Å model of the mutant protein (PDB ID: **5W4D**). We used this model for molecular replacement into the wild-type data set to build a complete 1.5 Å model (PDB ID: **5W4A**). Data and model statistics are in **Table S2**. Model coordinates and data are available at RCSB (www.rcsb.org).

### Size exclusion chromatography with multi-angle laser light scattering (SEC-MALS)

Molecular weights of *C. elegans* PGL-3 NtDD wild type and mutant recombinant protein were determined by conducting SEC-MALS experiments using Agilent Technologies 1260 LC HPLS system (Agilent Technologies, Santa Clara, CA) equipped with Dawn® Heleos™II 18-angle MALS light scattering detector, Optilab® T-rEX™ (refractometer with EXtended range) refractive index detector, WyattQELS™ quasi-elastic (dynamic) light scattering (QELS) detector and ASTRA software (all four from Wyatt Technology Europe GmbH, Dernbach, Germany). A total of 500 µL (1 mg/mL) of the samples in HNT buffer (20 mM HEPES pH 7.5, 100 mM NaCl, 0.5 mM TCEP pH 7.4) were injected and run on a Superdex 75 10/300 GL column (GE Healthcare) pre-equilibrated with the same buffer, at a flow rate of 0.5 mL/min at 20°C. Lysozyme (Sigma-Aldrich, St. Louis, MO) was used as a control.

### Mammalian cell culture maintenance, transfection and imaging

Full length PGL-1 was cloned into a pcDNA 3.1 vector (Thermo Fisher Scientific, Waltham, MA) with a C-terminal eGFP and OLLAS epitope linker. Mutations to PGL-1 were created using Gibson Assembly cloning ^59^. Chinese Hamster Ovary (CHO) cells (ATCC, Manassas, VA) were propagated according to distributor’s recommendations. Briefly, cells were grown in F-12K Medium (Thermo Fisher Scientific, Waltham, MA) with 10% fetal bovine serum (Gibco), and split with Trypsin 0.25% (Gibco) every 2-3 days. Cells were grown to 70% confluence and transfected with TransIT-CHO Transfection Kit (Mirus Bio LLC, Madison, WI). Transfected cells were split the following day and grown in Ibitreat 15 u-Slide 8 well slides (Ibidi, Madison, WI) overnight. Hoechst stain (Invitrogen, Carlsbad, CA) was added to wells prior to imaging by confocal microscopy for GFP and Hoechst fluorescence, and transmitted light. Well dilutions were chosen based on adequate cell spacing to discern each cell, and 25 fields of view were taken based on the highest concentration of GFP-positive cells. Experiments were repeated four times with similar results. During image collection, we observed a single example of a granule-like blob in the PGL-1::OLLAS::GFP K126E K129E. The cell appeared unhealthy, and thus the granule may be an artifact of cell death, but we included it in our study for completeness.

### Worm maintenance, CRISPR mutagenesis, fertility and imaging

Frozen strains:

N2 Bristol

JK5687: *pgl-1(q894)*[*PGL-1::SNAP*] *IV*

JK5902: *pgl-1(q975)*[*PGL-1(R123E)::SNAP*] *IV*

JK6158: *wago-1 q1087*[*WAGO-1::3xV5*]*; pgl-1(q894)*[*PGL-1::SNAP*] *IV*

JK6159: *wago-1 q1089*[*WAGO-1(null deletion)::3xV5*]*; pgl-1(q894)*[*PGL-1::SNAP*] *IV*

JK6157: *wago-1 q1087*[*WAGO-1::3xV5*]*; pgl-1(q975)*[*PGL-1(R123E)::SNAP*] *IV*

JK5898: *glh-1(q858)*[*GLH-1::3xMYC*] *I; pgl-1(q894)*[*PGL-1::SNAP*] *IV; pgl-3(q861)*[*PGL-3::3xV5*] *V*

JK5970: *qSi375*[*(mex-5 promoter::eGFP::linker::his-58::3xboxb::tbb-2 3’UTR) *weSi2*] *II; pgl-1(q894)*[*PGL-1::SNAP*] *IV*

JK5873: *qSi375*[*(mex-5 promoter::eGFP::linker::his-58::3xboxb::tbb-2 3’UTR) *weSi2*] *II; pgl-1(q994)*[*PGL-1::SNAP::λN22*]*/nT1*[*qIs51*]*(IV;V)*

JK5874: *qSi375*[*(mex-5 promoter::eGFP::linker::his-58::3xboxb::tbb-2 3’UTR) *weSi2*] *II; pgl-1(q994)*[*PGL-1::SNAP::λN22*]*/nT1*[*qIs51*]*(IV;V)*

JK6149: *qSi375*[*(mex-5 promoter::eGFP::linker::his-58::3xboxb::tbb-2 3’UTR) *weSi2*] *II; pgl-1(q994)*[*PGL-1::SNAP::λN22*]*/nT1*[*qIs51*]*(IV;V)*

JK6150: *qSi375*[*(mex-5 promoter::eGFP::linker::his-58::3xboxb::tbb-2 3’UTR) *weSi2*] *II; pgl-1(q994)*[*PGL-1:SNAP:λN22*]*/nT1*[*qIs51*]*(IV;V)*

JK6147: *wago-1 q1089*[*WAGO-1(null deletion)::3xV5*]*; qSi375*[*(mex-5 promoter::eGFP::linker::his-58::3xboxb::tbb-2 3’UTR) *weSi2*] *II; pgl-1(q994)*[*PGL-1::SNAP::λN22*]*/nT1*[*qIs51*]*(IV;V)*

JK6148: *wago-1 q1089*[*WAGO-1(null deletion)::3xV5*]*; qSi375*[*(mex-5 promoter::eGFP::linker::his-58::3xboxb::tbb-2 3’UTR) *weSi2*] *II; pgl-1(q994)*[*PGL-1::SNAP::λN22*]*/nT1*[*qIs51*]*(IV;V)*

JK6367: *qSi375*[*(mex-5 promoter::eGFP::linker::his-58::3xboxb::tbb-2 3’UTR) *weSi2*] *II*

JK6368: *wago-1 q1089*[*WAGO-1(null deletion)::3xV5*]*; qSi375*[*(mex-5 promoter::eGFP::linker::his-58::3xboxb::tbb-2 3’UTR) *weSi2*] *II*

Worm strains that could not be frozen:

1. *pgl-1(q960)[PGL-1(K126E K129E)::SNAP] IV*

2. *glh-1(q858)[GLH-1::3xMYC] I; pgl-1(q960)[PGL-1(K126E K129E)::SNAP] IV; pgl-3(q861)[PGL-3::3xV5] V*

3. *qSi375*[*(mex-5 promoter::eGFP::linker::his-58::3xboxb::tbb-2 3’UTR) *weSi2*] *II; pgl-1(q1053)*[*PGL-1(K126E K129E)::SNAP::λN22*]*/nT1*[*qIs51*]*(IV;V)*

*C. elegans* were maintained as previously reported ^60^. For CRISPR-Cas9 mutagenesis, a Cas9 protein co-conversion approach was used ^61^. Briefly, worms were injected with a target CRISPR-Cas9 RNA (crRNA) or a plasmid expressing a Cas9-scaffold with tandem target sequence RNA (sgRNA) to a gene of interest ^61^, a target crRNA to *dpy-10* or *unc-58*, a scaffolding tracrRNA (IDT), recombinant Cas9 protein ^62^, a *dpy-10/unc-58* repair DNA oligo that inserted a dominant mutation ^61^, and an epitope tag/missense mutant repair oligo or PCR product. See below for a Table of guide RNAs and repair templates used. F1s with the co-injection marker phenotype were additionally screened by a combination of PCR without or with restriction enzyme digest to identify those with the repair of interest. In JK5687, a SNAP tag ^30^ was inserted between PGL-1 amino acids G713 and G714 in N2 worms. A 3xMYC tag was added to the N-terminus of GLH-1 between G17 and F18. A 3xV5 tag was added in the C-terminal region of PGL-3 between residues G627 and S628. A 3xV5 tag was added to the C-terminus of WAGO-1 between residues E914 and A915. To generate the *wago-1* null allele, a WAGO-1::3xV5 allele was mutated so that 648 base pairs were deleted from the N-terminus and proper coding frame shifted to add premature stop codons (**Figure S7C**). The *wago-1* null allele was confirmed by staining and imaging (**Figure 6D**). F2s were PCR screened to identify homozygous SNAP alleles and the PCR product sequenced to confirm proper repair. Three worm strains were too infertile to freeze. All worms were outcrossed at least twice with N2, with the exception of *glh-1(q858)*[*GLH-1::3xMYC*] *I; pgl-1(q960)*[*PGL-1(K126E K129E)::SNAP*] *IV; pgl-3(q861)*[*PGL-3::3xV5*] *V* that was backcrossed with JK5898.

Worms were singled into the peripheral wells of a 24-well plate that contained NGM agar and OP50 bacteria. Worms were allowed to propagate for 5 days at 20°C or 25°C, and then scored for progeny and gravid progeny. We report the progeny numbers here.

### CRISPR-Cas9 guide RNAs and repair oligos

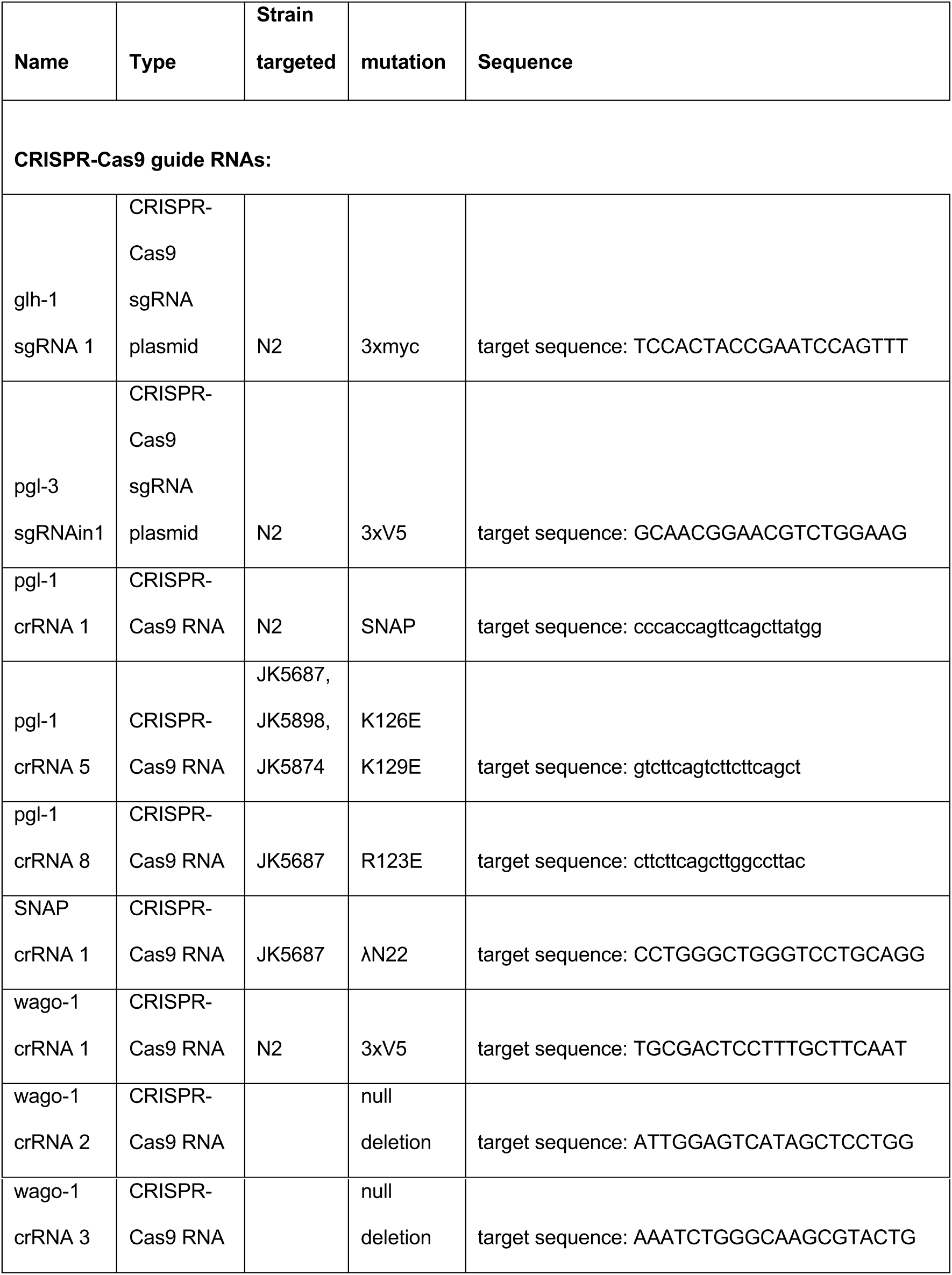

### CRISPR-Cas9 repair oligos

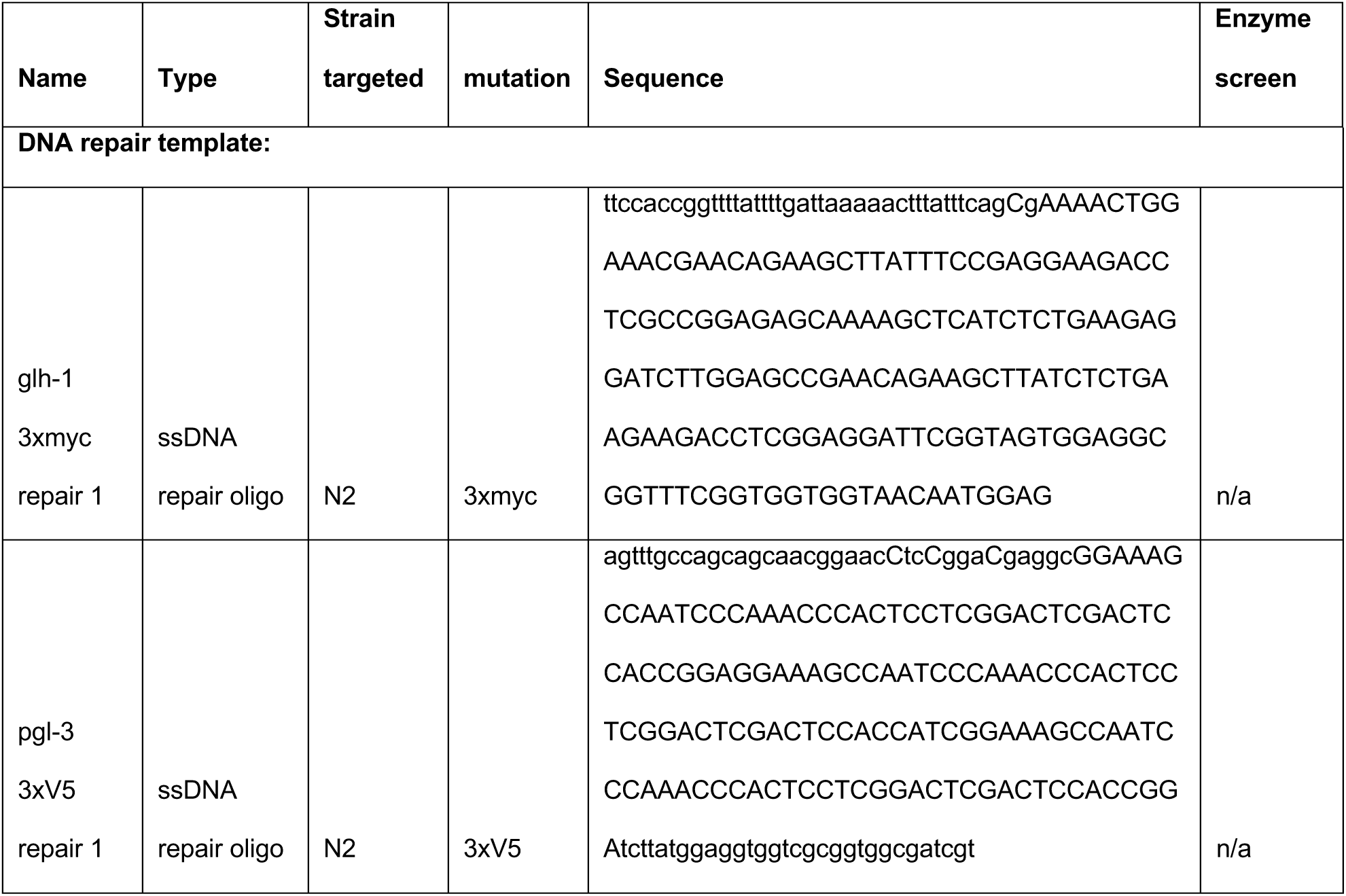
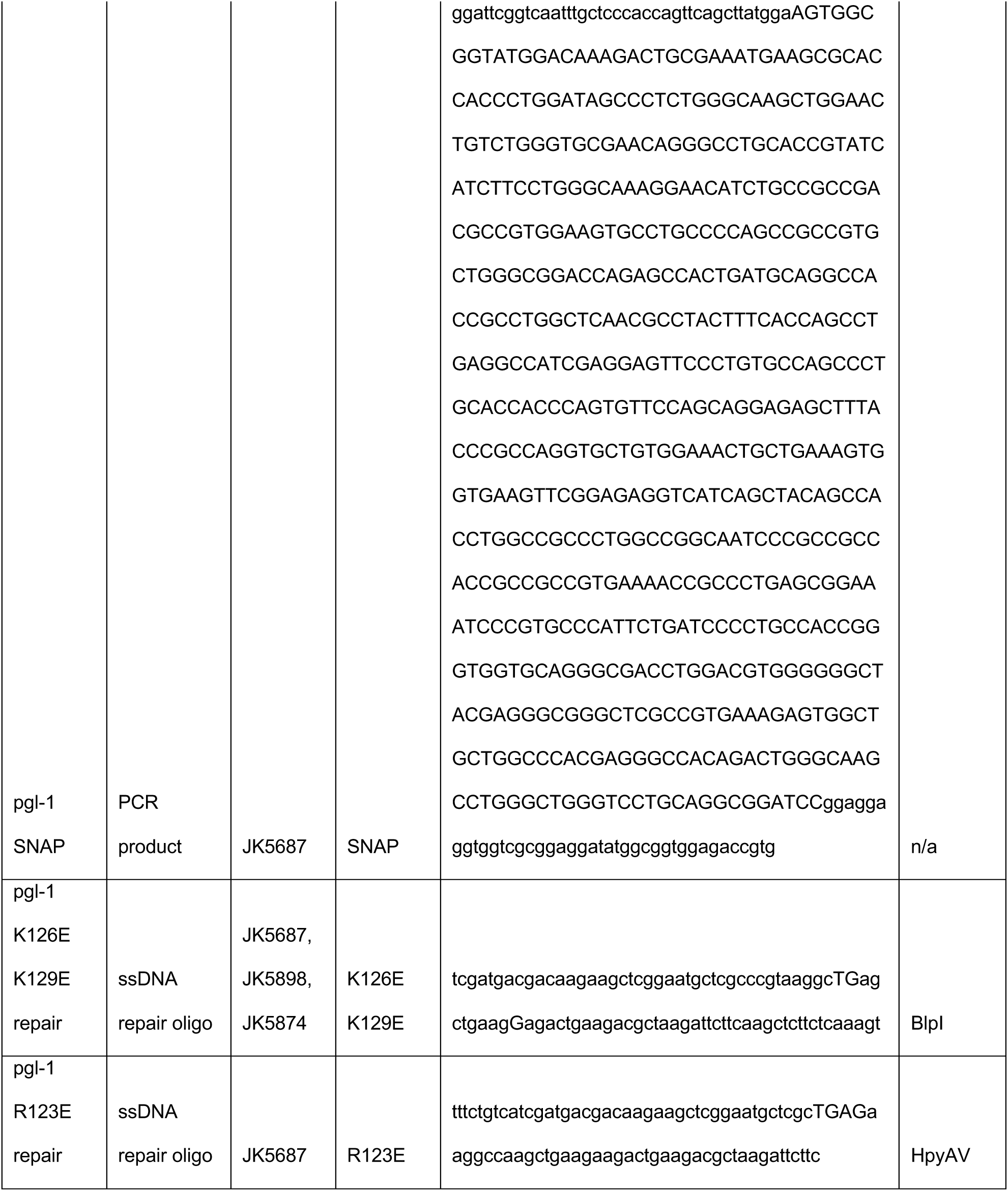
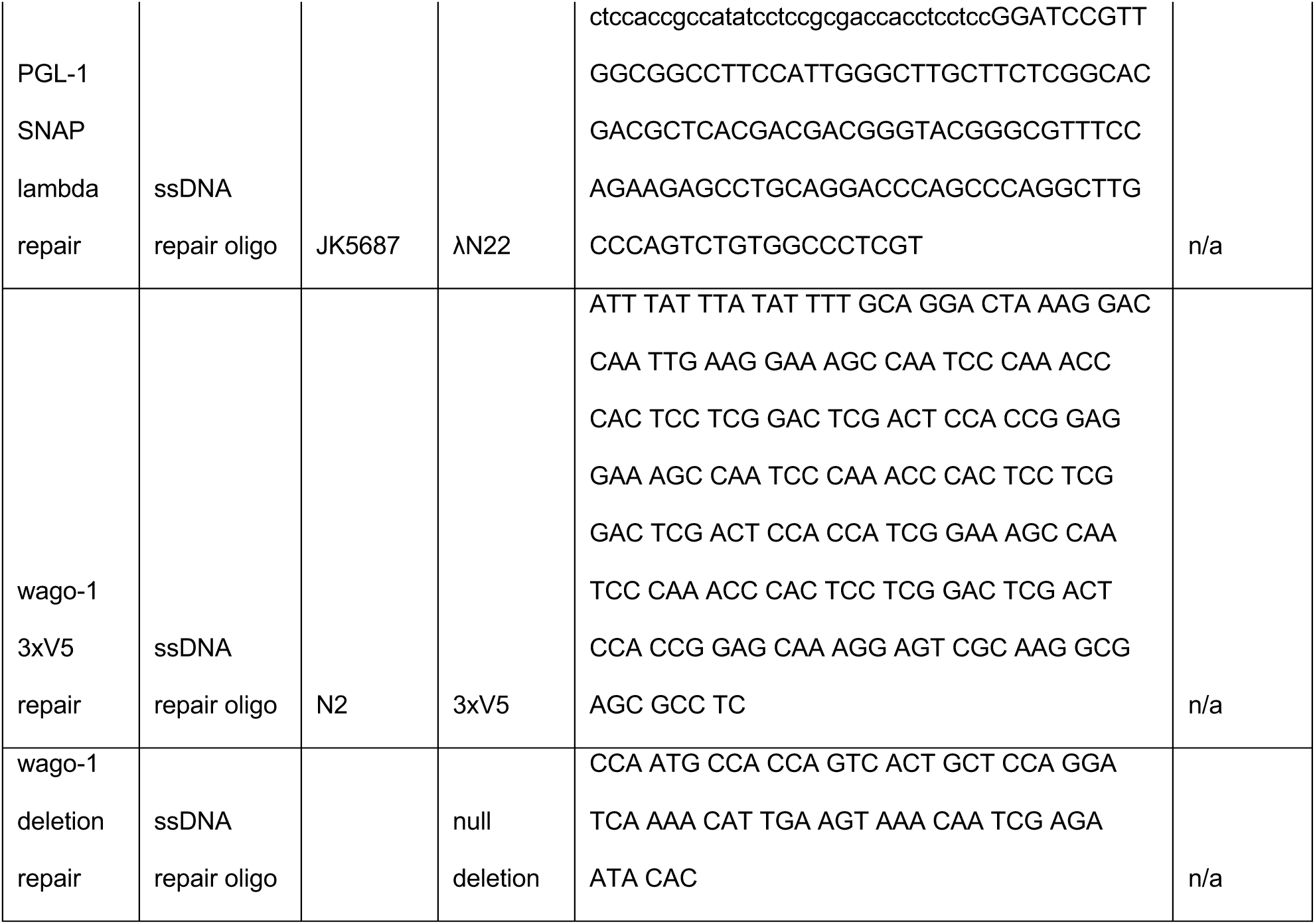

### Immunoblot

To check PGL-1::SNAP protein expression, worms were boiled in 5x sample buffer (250 mM Tris pH 6.8, 25 mM EDTA pH 8.0, 25% glycerol, 5% SDS, 500 mM beta-mercaptoethanol), run on an SDS-PAGE gel, and transferred to PVDF. The blot was blocked in 5% dehydrated milk in PBS-T, incubated overnight with anti-SNAP antibody (NEB, Ipswich, MA), washed and probed with goat-anti-rabbit horseradish peroxidase secondary antibody (Invitrogen, Carlsbad, CA), and developed with ECL substrate (Thermo Fisher Scientific, Waltham, MA) and film (Kodak, Rochester, NY). Radiographs were scanned, contrast adjusted and cropped (Photoshop, Adobe Creative Cloud) as shown.

### Fluorescent imaging

To analyze GFP reporter expression, L4 larvae were propagated for approximately 24 hours to adulthood at 20°C, placed in M9 with 0.1 mM levamisole on a glass slide with a cover slip, imaged at 10x magnification on a compound microscope and counted for the presence or absence of GFP fluorescence in their germlines. Numbers represent totals from two separate experiments. The reporter images of live worms were taken of worms treated in a similar manner and visualized on a Leica SP8 scanning laser confocal microscope.

For confocal imaging, germlines were extruded, fixed with 1% paraformaldehyde (Electron Microscopy Sciences, Hatfield, PA) and permeabilized with 0.5% Triton-X as previously described ^63^. Germlines were incubated with 1 μg/ml primary antibodies overnight [anti-MYC (JAC6 (rat), Bio-Rad, Hercules, CA); anti-V5 (sv5-Pk1 (mouse), Bio-Rad, Hercules, CA)] or 30 nM SNAP JF 549 ligand ^64^ for 1 hour, stained with fluorophore-labeled secondary antibodies (Alexa 488 Donkey anti-Mouse, Alexa 555 Donkey anti-Mouse, Alexa 647 Donkey anti-Mouse, Alexa 488 Goat anti-Rabbit; Invitrogen, Carlsbad, CA) and DAPI (Invitrogen, Carlsbad, CA), washed and mounted in Vectashield (Vector Laboratories, Burlingame, CA). Quantitation of GFP and SNAP fluorescence was performed in ImageJ. Graph made in Excel. Quantitation of GFP and SNAP fluorescence were quantitated in summed intensity projections of confocal stacks in ImageJ, similar to Crittenden *et al.* ^65^. Briefly, a line (40 pixel width) was drawn along the germline axis. Pixel intensity was measured using plot profile, and intensities averaged and plotted in Excel.

### Single molecule fluorescence *in situ* hybrididation (smFISH)

For smFISH, gonads were extruded, fixed, and hybridized with single molecule FISH probes as described ^31^. The *gfp* exon probe set contains 38 unique oligonucleotides labeled with CAL Fluor Red 610. Briefly, probes were dissolved in RNase-free TE buffer (10 mM Tris-HCl, 1 mM EDTA, pH 8.0) to create a 250 µM probe stock. Mid-L4 stage animals were grown on OP50 for 24 hours, then dissected in PBS+0.1% Tween-20 + 0.25 mM levamisole. Animals were fixed in 4% paraformaldehyde for 20 minutes, incubated at room temperature in PBS-T (PBS + 0.1% Tween-20) for 10-25 minutes, and equilibrated in smFISH wash buffer (30 mM sodium citrate pH 7.0, 300 mM NaCl, 1% formamide, 0.1% Tween-20, DEPC water) for 10-16 minutes. Samples were then incubated in hybridization buffer (30 mM sodium citrate pH 7.0, 300 mM NaCl, 1% formamide, 10% dextran sulfate w/v, DEPC water) plus 0.5 µM smFISH probe at 37°C for 26-44 hours. 30 nM SNAP 549 ligand was added during the smFISH wash buffer + DAPI wash; samples were washed at 37°C for approximately 60 minutes. Finally, samples were resuspended in 12 µL Antifade Prolong Gold mounting medium (Thermo Fisher Scientific, Waltham, MA), mounted on glass slides, and cured in a dark drawer for at least 24 hours before imaging.

### Confocal imaging for smFISH and protein fluorescence

Samples were imaged using a Leica SP8 scanning laser confocal microscope, taking 0.3 µm (smFISH experiments) or 1 µm (other non-smFISH confocal imaging experiments) slices in sequence. Maximum intensity partial stack projections were generated and brightness adjusted using ImageJ ^66^. All images were treated equally in ImageJ and Photoshop, with the exception of the transmitted light images. Imaging experiments were repeated at least twice with similar results, with the exception of PGL-1(K126E K129E)::SNAP::λN22 worms.

### smFISH image analysis

To quantify *gfp* smFISH signal localization between samples, *gfp* smFISH signal and PGL-1::SNAP signal were each detected then colocalized using Imaris version 9.3.1. The same batch settings were run on all directly-compared images. In the 3D view for each image, detection for both *gfp* and PGL-1::SNAP surfaces was checked. If necessary, the edit tool was used to select and delete *gfp* and/or PGL-1::SNAP surfaces that were detected outside the imaged germline (e.g. on nearby intestine tissue). The *gfp* surfaces object was selected and the total *gfp* signal was pulled from the statistics tab (Detailed tab, select Average Values from dropdown, “Intensity Sum Ch=4 Img=1” row and “Sum” column) and stored in an excel file. The Surface-surface coloc was run from Surface-Surface coloc XTension. The *gfp* and PGL-1::SNAP channels were selected and the “no smoothing” option was selected. The same statistics tab navigation was used to pull GFP intensity from the ColocSurface objects that were created by the XTension. Total GFP intensity in the colocalized surfaces was divided by total GFP intensity in the germline to calculate how much GFP signal is located in the PGL granules. Some *gfp* and PGL::SNAP co-localization values were calculated as over 100%. Although colocalization was performed using the Imaris user interface, the Surface-Surface Colocalization XTension ran an available MATLAB algorithm. We observed that the Surface-Surface colocalization created a new set of colocalization surfaces, which are differently shaped and larger than the surfaces written with Imaris algorithms to detect *gfp* or PGL:SNAP surfaces. Therefore, the total *gfp* intensity in the coloc surfaces was sometimes larger than in the *gfp* surfaces.

PGL granule staining is sensitive to physical perturbation and some germlines lacked PGL-1::SNAP signal entirely or almost entirely. GFP smFISH and PGL-1::SNAP signal overlap was therefore calculated from the top 66% of images in terms of PGL-1::SNAP Intensity (Detailed tab, select Average Values from dropdown, “Intensity Sum Ch=5 Img=1” row and “Sum” column). Excluding the low PGL images increased median signal overlap from 26% to 34% in PGL-1::SNAP and from 41% to 57% in PGL-1::SNAP::λN22 (**Figure S2O**). Excluding the low PGL images increased median signal overlap from 113% to 121% in PGL-1::SNAP::λN22 with WAGO-1 and from 74% to 81% in PGL-1::SNAP::λN22 without WAGO-1 (**Figure S9I**).

Each of the two independent samples were run through a two-tailed t test to calculate significance. We used the online calculator at http://vassarstats.net.

### Imaris batch detection settings

|  | GFP smFISH<br>channel: Figure<br>S2L | PGL:SNAP<br>channel:<br>Figure S2L | GFP smFISH<br>channel:<br>Figure S9I | PGL:SNAP<br>channel:<br>Figure S9I |
| --- | --- | --- | --- | --- |
| [Algorithm] |  |  |  |  |
| Enable Region of<br>Interest | false | false | false | false |
| Enable Region<br>Growing | true | true | true | true |
| Enable Tracking | false | false | false | false |
| [Source Channel] |  |  |  |  |
| Source Channel Index | 4 | 5 | 4 | 5 |
| Enable Smooth | true | true | true | true |
| Surface Grain Size | 0.090233<br>(default -- 2x<br>pixel size) | 0.090233<br>(default -- 2x<br>pixel size) | 0.090241<br>(default) | 0.090241<br>(default) |
| Enable Eliminate Background | true | true | true | true |
| Diameter of largest sphere | 0.28000 um* | 0.60000 um* | 0.28000 um | 0.60000 um |
| [Threshold] |  |  |  |  |
| Enable Automatic Threshold | false | false | false | false |
| Manual Threshold value | 2.0241** | 5.66575** | 3.08326 | 15.9772 |
| Active Threshold | true | true | true | true |
| Enable Automatic Threshold B | true | true | true | true |
| Manual Threshold Value B | 19.1165** | 14.5472** | 97.9303 | 67.4159 |
| Active Threshold B | false** | false** | false | false |
| Region Growing Estimated Diameter | 0.30000 um* | 0.40000 um* | 0.30000 um | 0.40000 um |
| [Classify Seed Points] |  |  |  |  |
| "Quality" | above 1.2526* | above 1.9952* | above 3.5559 | above 15.052 |
| [Classify Surfaces] |  |  |  |  |
| "Number of Voxels Img = 1" | above 28.000* | above 28.000* | above 28.000 | above 28.000 |

\*Values were determined by trial and error, using values from measurements where possible.
\*\*Manual threshold initially set by turning on Upper Threshold (threshold B) -- this changes the value for the automatic lower threshold. Set the lower threshold to lock that value, then turn off the upper threshold. This makes the Value B stored, although it isn't actually in use (active = false). Note: the same technique would provide different values for each image, so the 2.0241 was arbitrarily set on the image settings.

We used a Dell Precision 5820 with a 64-bit Windows 10 Education operating system, an Intel(R) Xeon(R) W-1245 CPU @3.70GHz processor, and 128 GB of RAM. Imarisx64 9.3.1 and ImarisFileConverterx64 9.3.1 were installed along with MATLAB R2018b. The Surface-Surface Colocalization XTension was downloaded and installed from the Bitplane XTension File Exchange at http://open.bitplane.com/tabid/235/Default.aspx?id=111.

## Acknowledgements

The authors thank M. Cox for equipment; J. Claycomb for plasmids and strains; L. Lavis for supplying the SNAP JF 549 ligand; T. Hoang, S. Strome, D. Updike, and members of the Kimble and Wickens labs for helpful discussions. Use of the LS-CAT Sector 21 was supported by the Michigan Economic Development Corporation and the Michigan Technology Tri-Corridor (Grant 085P1000817). The Keck Biophysics Facility (SEC-MALS) is supported in part by NCI CCSG P30 CA060553 grant awarded to the Robert H Lurie Comprehensive Cancer Center. STA was supported by K99HD081208. CAB was supported by NIH grants GM094584, GM094622 and GM098248. MW was supported by NIH grant GM50942. JK is an Investigator of the Howard Hughes Medical Institute.

## Author contributions

STA conceived and performed experiments, analyzed data and wrote the paper. TRL and SLC performed experiments, analyzed data and helped write the paper. CAB performed experiments and analyzed data. MW analyzed data and helped write the paper. JK conceived experiments, analyzed data and wrote the paper.

## Competing interests

The authors declare no competing interests.

## Supplemental Tables and Figures

**Figure S1.**
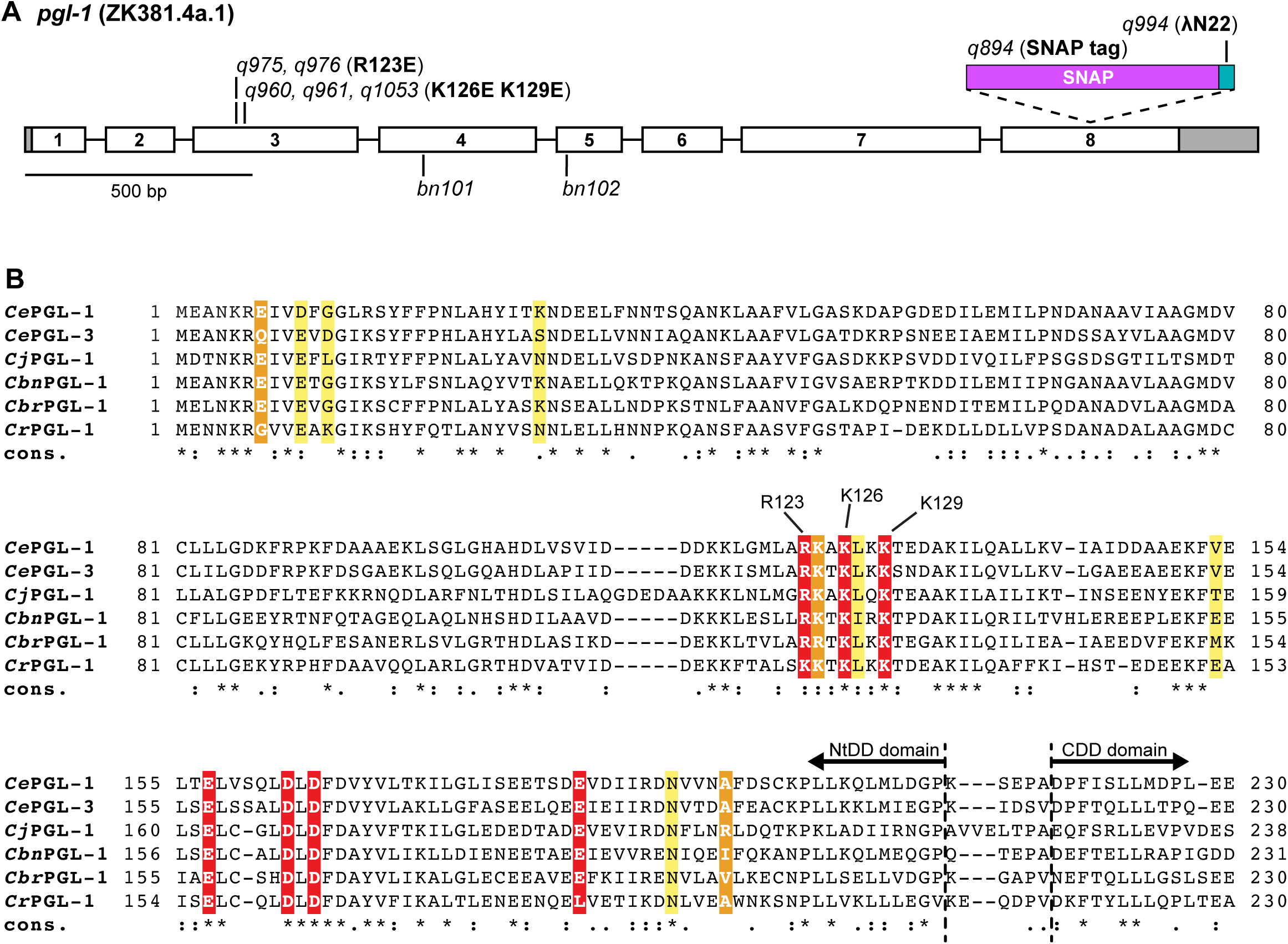
PGL sequence alignment and the *pgl-1* locus (A) *pgl-1* primary transcript. 5’ and 3’ UTRs are grey, exons are white, numbered 1-8 and separated by introns. Sites of *pgl-1* mutations are labeled, including location of SNAP tag (magenta) and λN22 fusion (blue). (B) Sequence alignment of PGL NtDD domain in *C. elegans* (*Ce*), *C. japonica* (*Cj*), *C. brenneri* (*Cbn*), *C. briggsae* (*Cbr*), *C. remanei* (*Cr*). Alignment and conservation (cons.) determined by T-Coffee ^67^. Starred residues (*) are identical. Period (.) and colon (:) residues are similar. Residues participating in salt bridges only are in orange. Residues participating in hydrogen bonds only are in yellow. Residues forming both hydrogen bonds and salt bridges are highlighted in red. *C. elegans* PGL-1 missense mutations and their allele numbers are labeled. Dashed lines mark the newly-annotated end of PGL-1 NtDD domain and start of PGL-1 CDD domain.

**Figure S2.**
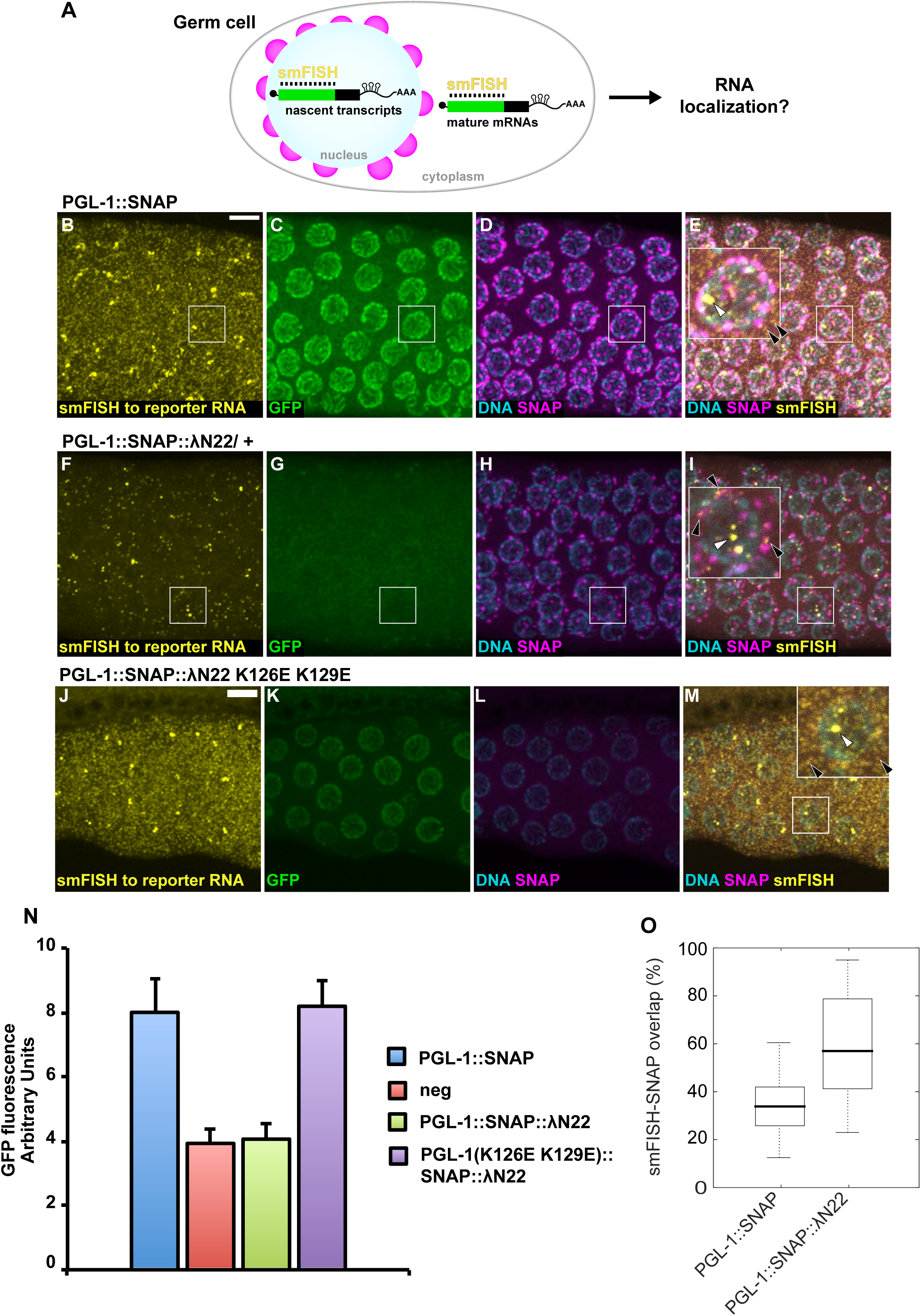
GFP reporter transcripts localize to P-granules when tethered (A) Single molecule fluorescent *in situ* hybridization (smFISH) in nematode germ cells to visualize RNA expression and localization. (B-M) Gonads were extruded from animals harboring the *gfp* reporter and (B-E) PGL-1::SNAP (n=30); (F-I) PGL-1::SNAP::λN22 (n=27); (J-M) PGL-1 (K126E K129E)::SNAP::λN22 (n=10). Gonads were fixed and imaged for *gfp* RNA using (B,F,J) smFISH; (C,G,K) GFP protein fluorescence; (D,H,L) DNA (DAPI) and SNAP. The smFISH, DNA and SNAP images are merged in E,I,M. White arrows mark examples of intranuclear puncta; black arrows mark examples of cytoplasmic puncta. Scale bar, 5 µm, for all images, except for 2.5-fold enlarged images in inset. For germline location, see **Figure S5A**. (N) Quantification of GFP signal in confocal images for the GFP reporter expressed with PGL-1::SNAP (n=23), PGL-1::SNAP::λN22 (n=24), PGL-1 (K126E K129E)::SNAP::λN22 (n=10). A silenced GFP reporter (n=25) served as a negative control. Mean GFP fluorescent signal and standard deviation reported in Arbitrary Units. (O) smFISH and PGL-1::SNAP signal colocalization in germlines expressing the GFP reporter and either PGL-1::SNAP or PGL-1::SNAP::λN22. Reported as the (Sum of smFISH GFP Intensity in SNAP:smFISH colocalized binned signal)/(Total smFISH GFP binned signal). Box plots represent the first and third quartiles, black line is median, whiskers to min and max values. Co-localization greater in germlines expressing PGL-1::SNAP::λN22 (*p*<0.01). See Methods for more details.

**Figure S3.**
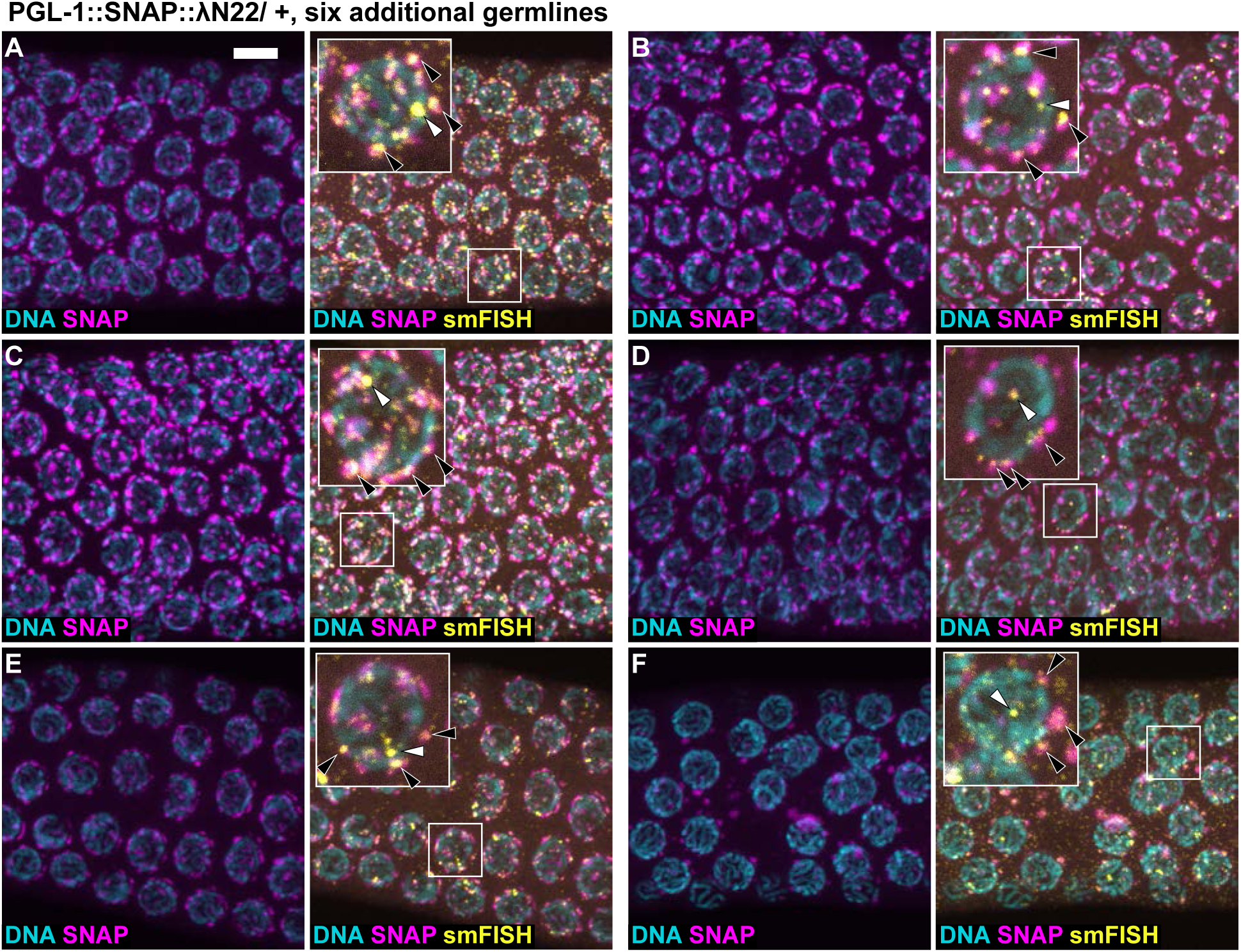
GFP reporter transcripts localize to P-granules when tethered (A-F) Six additional examples of germlines harboring PGL-1::SNAP::λN22 (n=27). Gonads were fixed and imaged for *gfp* RNA using smFISH, DNA (DAPI) and SNAP. The three are merged in images on right. White arrows mark examples of intranuclear puncta; black arrows mark examples of cytoplasmic puncta. Scale bar, 5 µm, for all images, except for 2.5-fold enlarged images in inset. For germline location, see **Figure S5A**.

**Figure S4.**
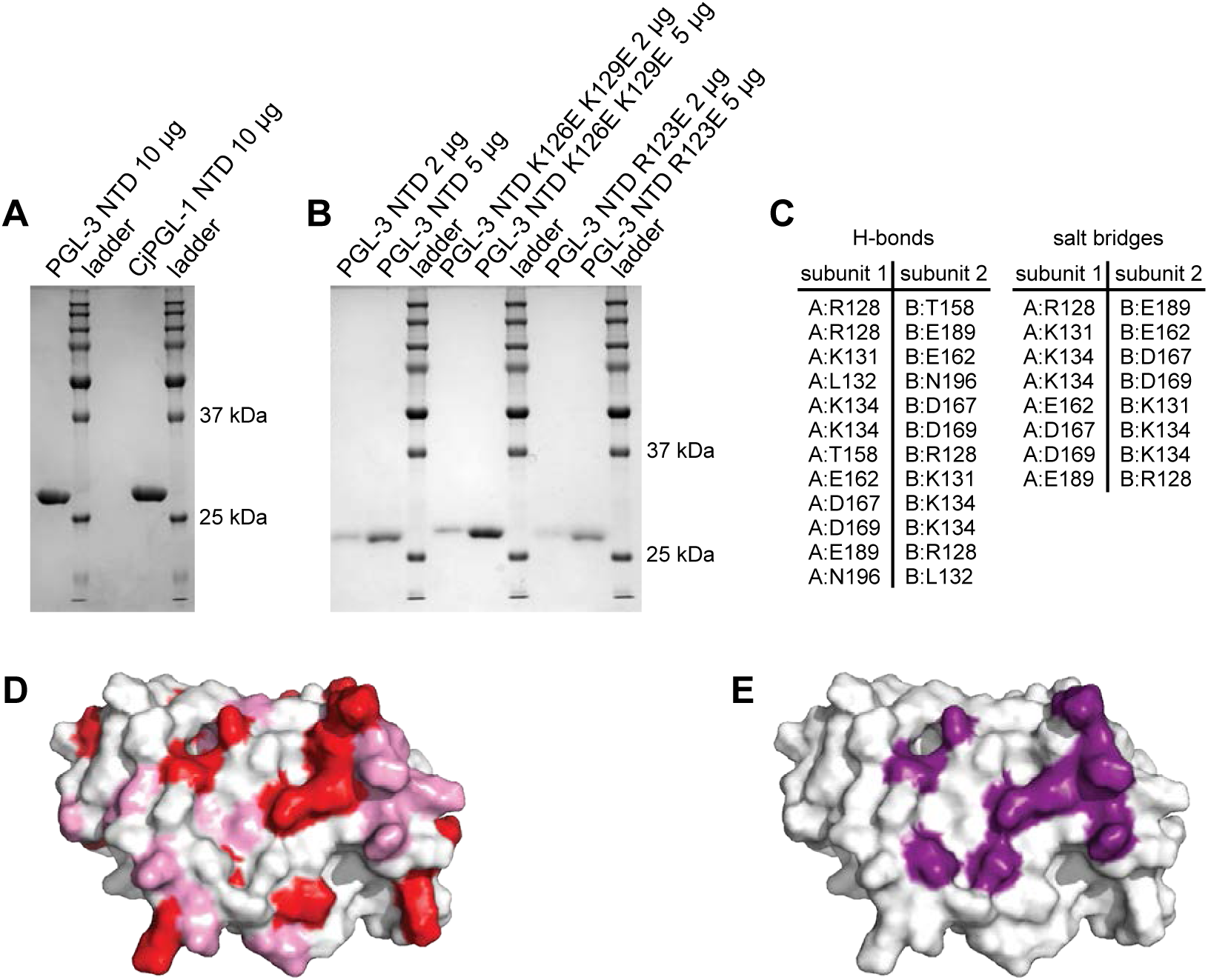
Supplemental biochemical and structural analyses of PGL NtDD (A,B) Coomassie-stained polyacrylamide gel of recombinant PGL NtDD wild-type and mutant protein. Ladder marker sizes labeled in kilodaltons (kDa) on right. (A) Recombinant *C. japonica* (Cj) PGL-1 NtDD protein used for crystallization. Recombinant *C. elegans* PGL-3 NTD protein included for comparison. (B) Wild-type and mutant *C. elegans* PGL-3 NtDD recombinant proteins used for biochemical characterization. (C) Tables of predicted hydrogen bonds and salt bridges at the NtDD dimerization interface. Amino acid numbers correspond to *C. japonica* PGL-1 NtDD. (D,E) Surface representation of the NtDD dimerization interface. (D) Amino acids colored by identity (red) and similarity (pink). (E) Amino acids at dimerization interface (purple).

**Figure S5.**
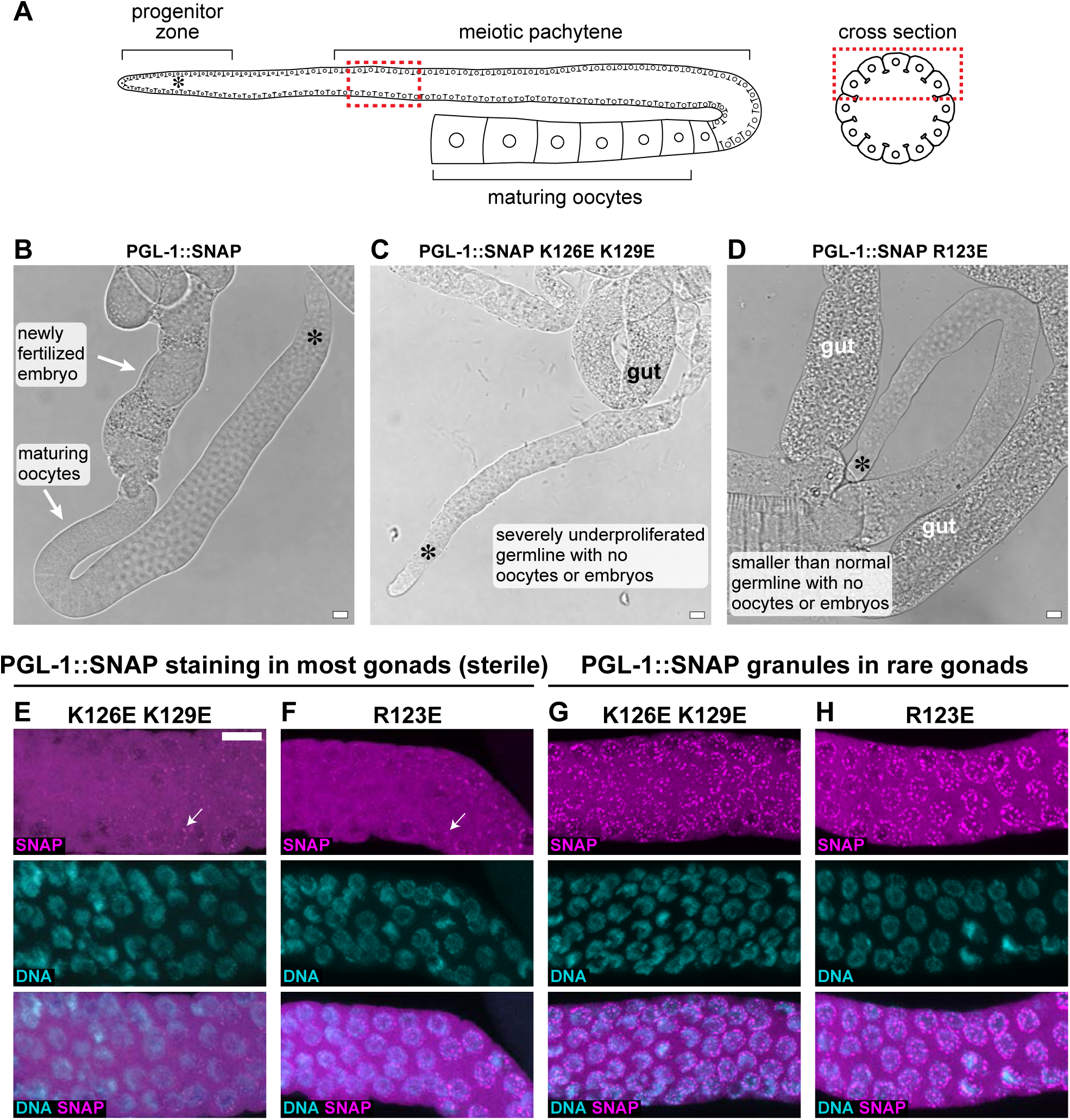
Supplemental images of PGL-1 dimerization mutants (A) Schematic of an adult hermaphrodite germline. An asterisk marks proliferating germ cells here and in images B-D. The germline produces oocytes at this stage; sperm were made earlier and stored in the spermatheca (not shown). Red box marks region imaged. (B-D) Representative brightfield images of extruded gonads from worms grown at 20°C. Scale bar, 10 μm. (B) Wild-type PGL-1::SNAP gonads are of normal size and produce oocytes and embryos. (C) Representative image of PGL-1(K126E K129E)::SNAP sterile gonads, which are small and produce no gametes. (D) PGL-1(R123E)::SNAP sterile gonads are also small and produce no gametes. (E-H) Representative partial z-projection stacks of SNAP and DNA stained germlines. Scale bar, 10 μm, applies for all images. (E,F) PGL-1::SNAP mutant protein is expressed in sterile gonads (no embryos observed). (E) PGL-1(K126E K129E)::SNAP, n=21 gonads imaged. (F) PGL-1(R123E)::SNAP, n=30 gonads imaged. (G,H) In single rare gonads, PGL-1::SNAP mutants were seen to assemble into granules in all germ cells.

**Figure S6.**
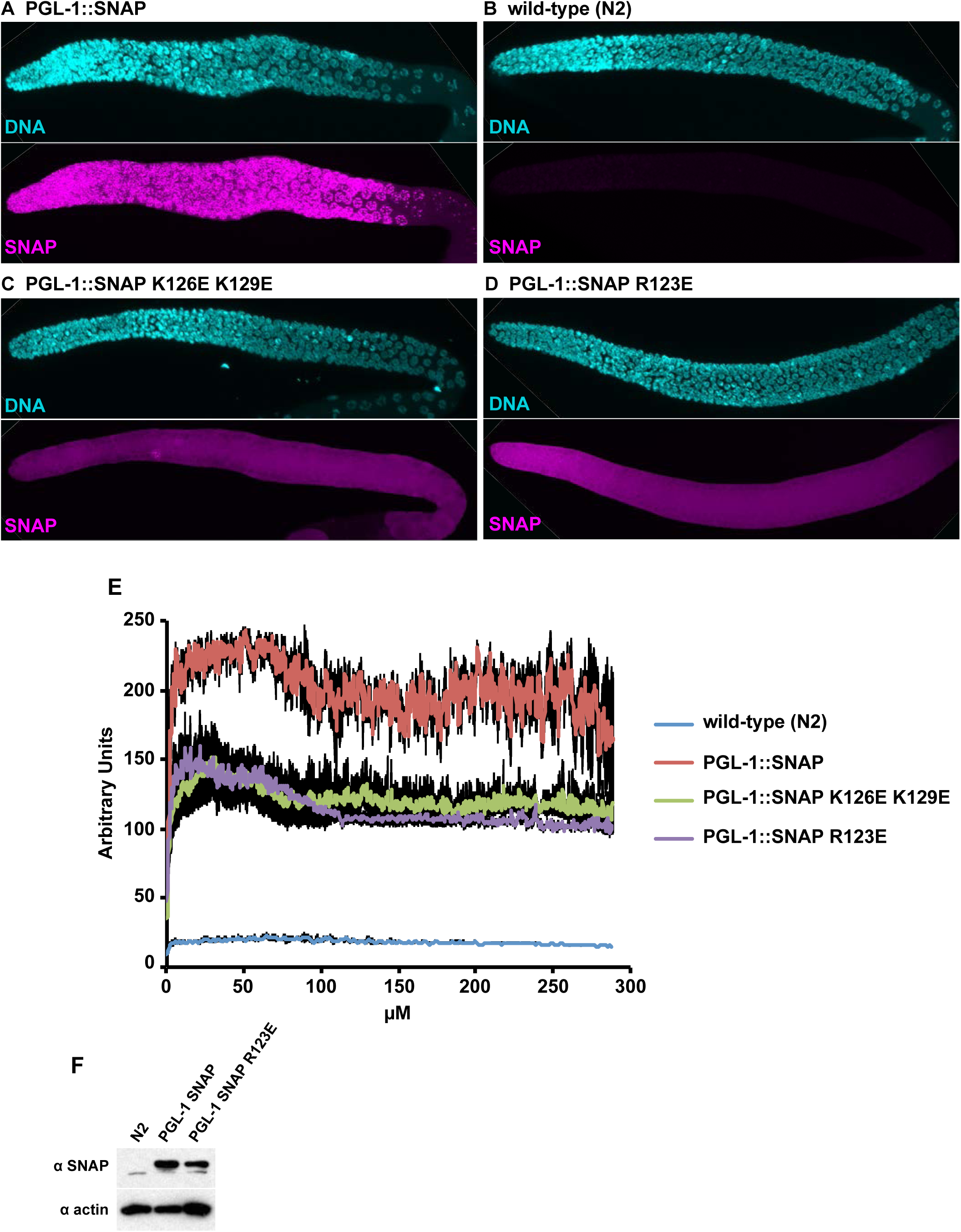
PGL-1 protein expression. (A-D) Examples of SNAP staining in (A) PGL-1::SNAP, (B) wild-type N2 negative control, (C) PGL-1(K126E K129E)::SNAP and (D) PGL-1(R123E)::SNAP. (E) Fluorescent signal (y-axis, arbitrary units) from SNAP-stained PGL-1::SNAP, N2, PGL-1(K126E K129E)::SNAP and PGL-1(R123E)::SNAP germlines were measured from the beginning of the germline (start of progenitor zone, **Figure S5A**) to the meiotic pachytene region (x-axis, μM). Solid line represents the mean with black showing the standard error. (F) Immunoblot of embryo-containing adult, N2, PGL-1::SNAP or PGL-1(R123E)::SNAP expressing worms. Samples were separated by SDS-PAGE and probed with SNAP and actin antibodies. Experiment was repeated twice with similar results.

**Figure S7.**
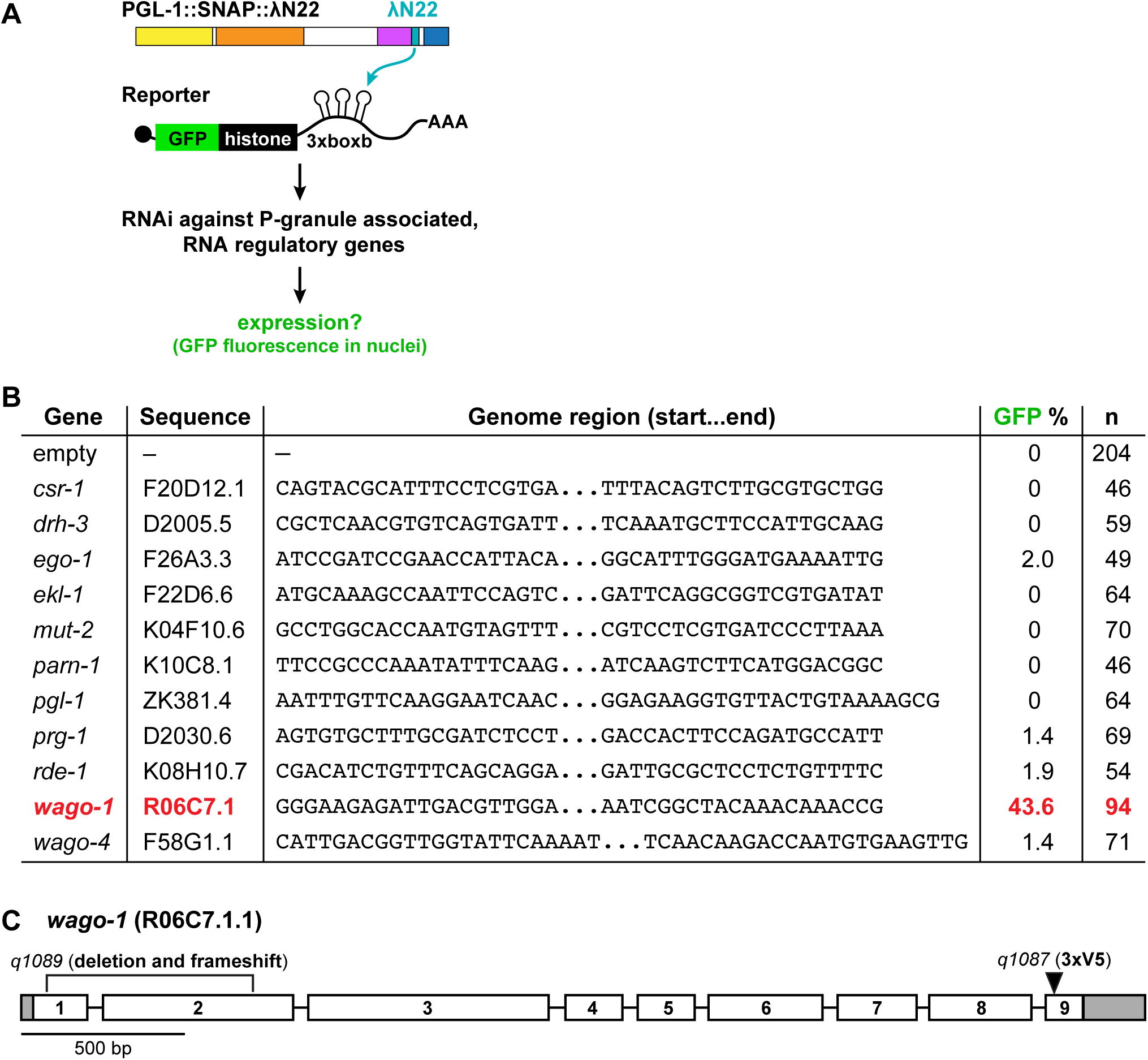
Targeted RNAi screen reveals WAGO-1 as a potential candidate in repressing PGL-tethered mRNA transcripts (A) Candidate gene RNAi screening strategy. PGL-1::SNAP::λN22 tethered to GFP reporter mRNAs repress GFP expression. We propagated larval worms on bacteria expressing RNAi for specific P-granule-associated enzymatic factors. After 5 days (one generation), young adult worm progeny were observed for GFP nuclei in their germlines, indicating tethering de-repression. (B) Candidate genes and their effect on GFP expression. Sequences of candidate genomic regions reported. Results were repeated at least three times and summarized here. (C) *wago-1* (R06C7.1.1) primary transcript. 5’ and 3’ UTRs are grey, exons are white, numbered 1-9 and separated by introns. Site of 3xV5 insertion and deleted region of null allele noted.

**Figure S8.**
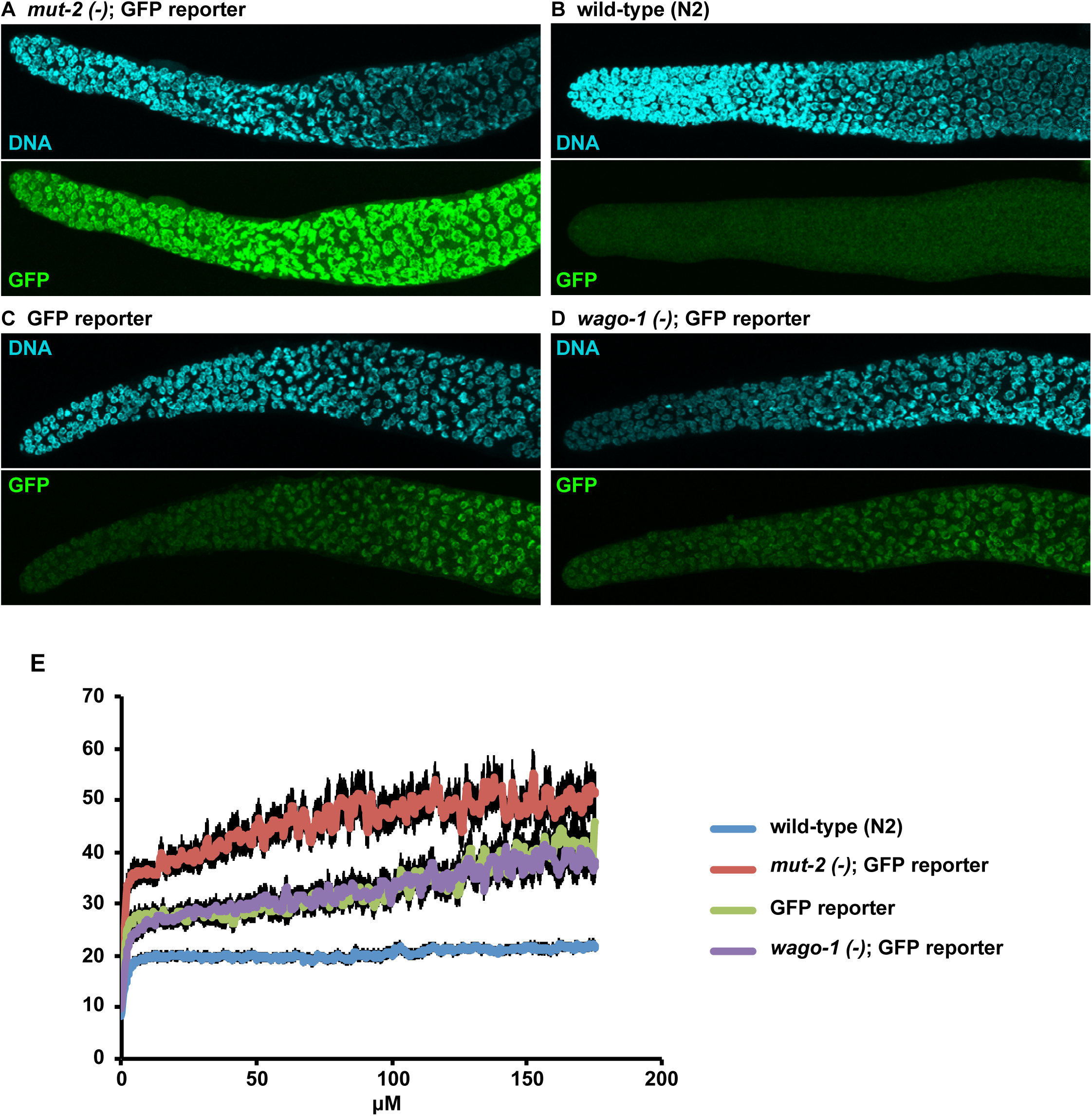
GFP reporter expression in the presence or absence of WAGO-1. (A-D) Examples of GFP fluorescence in (A) positive control *mut-2(-)*; GFP reporter, (B) negative control wild-type N2, (C) GFP reporter, and (D) *wago-1(-)*; GFP reporter. *mut-2* prevents reporter silencing. (E) GFP Fluorescent signal (y-axis, arbitrary units) from germlines of described genotypes were measured from the beginning of the germline (start of progenitor zone, **Figure S5A**) to the meiotic pachytene region (x-axis, μM). The greater than 100% values are due to the use of both Imaris and Matlab for quantitation (see Methods). Solid line represents the mean with black showing the standard error. Note that the GFP reporter expressed similarly with or without WAGO-1.

**Figure S9.**
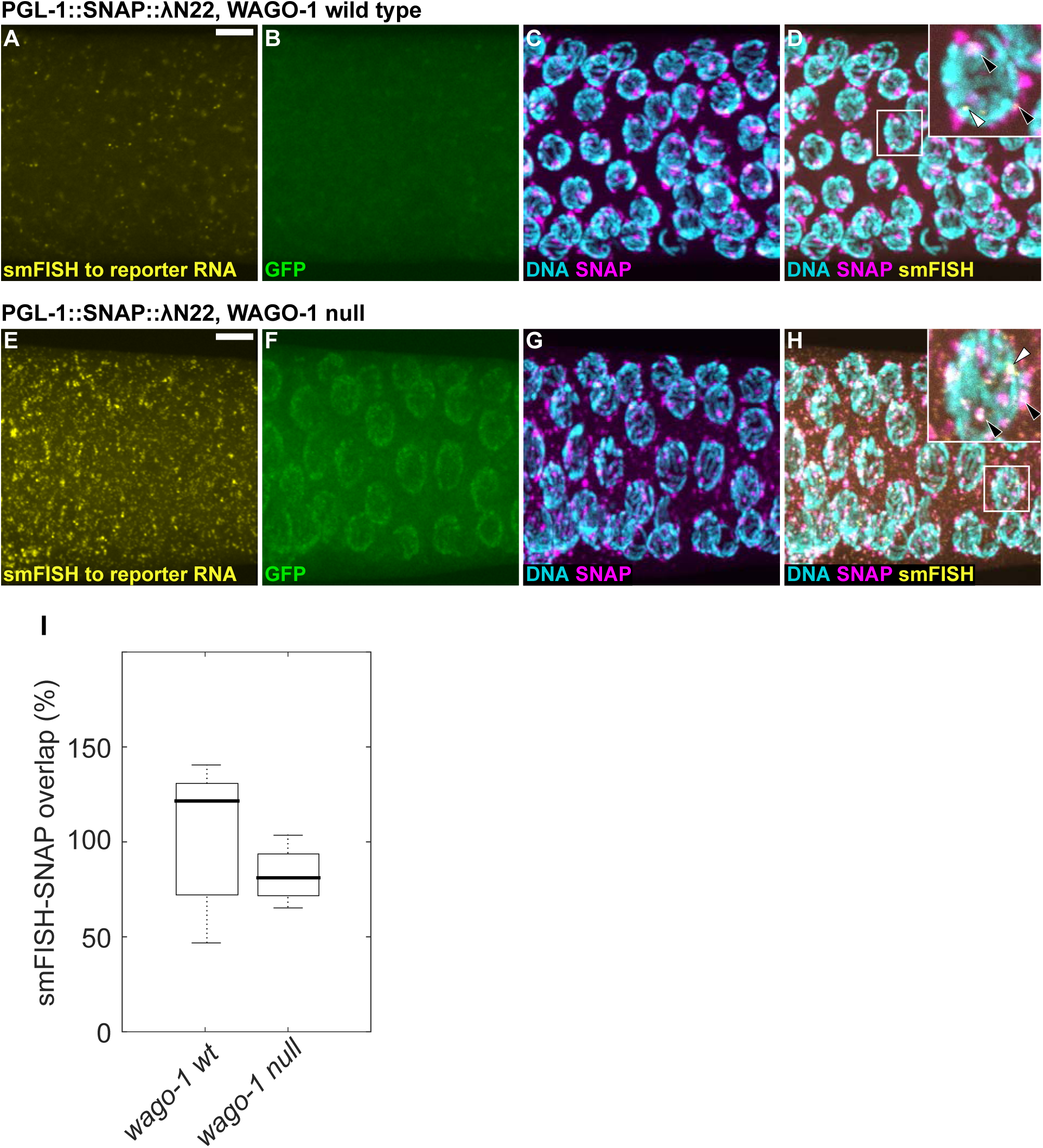
PGL-1-tethered GFP reporter transcripts localize to granules in the presence or absence of WAGO-1 Gonads were extruded from *gfp* reporter, PGL-1::SNAP::λN22 animals harboring (A-D) *wago-1* wild-type (n=22); (E-H) *wago-1* null (n=21). Gonads were fixed and imaged for (A,E) *gfp* RNA using smFISH, (B,F) GFP protein fluorescence, (C,G) DNA (DAPI) and SNAP. *gfp* RNA smFISH, DNA and SNAP merged in D, H. White arrows mark examples of intranuclear puncta; black arrows mark examples of cytoplasmic puncta. Scale bar, 5 µm, for all images, except for 2.5-fold enlarged images in inset. For germline location, see **Figure S5A**. (I) smFISH and PGL-1::SNAP signal colocalization in germlines expressing the GFP reporter, PGL-1::SNAP::λN22 in the presence or absence of WAGO-1. Reported as the (Sum of smFISH GFP Intensity in SNAP:smFISH colocalized binned signal)/(Total smFISH GFP binned signal). Box plots represent the first and third quartiles, black line is median, whiskers to min and max values. Co-localization similar in germlines expressing PGL-1::SNAP::λN22 with or without WAGO-1 (p=0.039). See Methods for more details.

**Figure S10.**
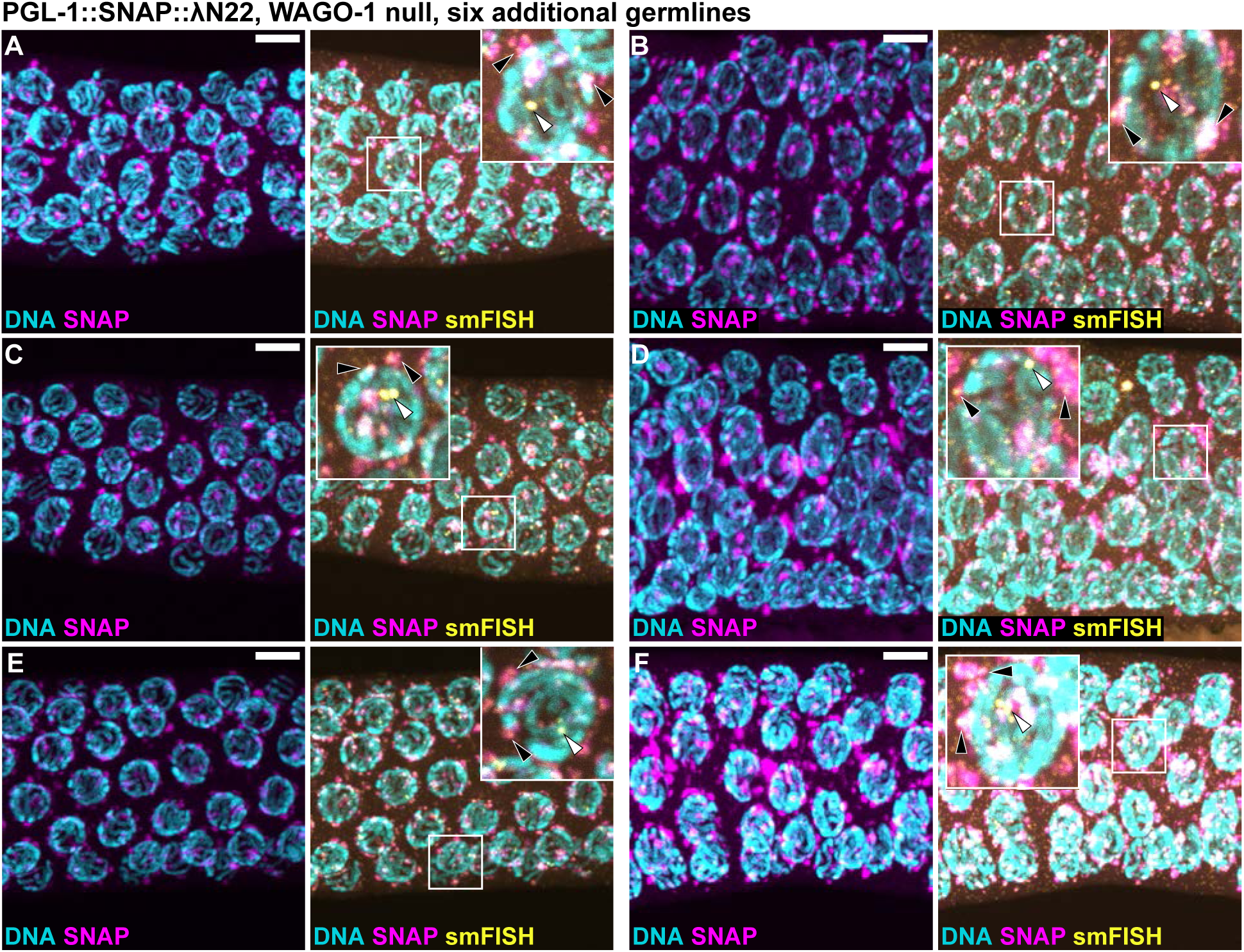
Additional images of PGL-1-tethered GFP reporter transcripts without WAGO-1 (A-F) Six additional examples of germlines harboring PGL-1::SNAP::λN22, *wago-1* null imaged for *gfp* RNA, DNA, and SNAP. *gfp* RNA, DNA and SNAP merged in right images. White arrows mark examples of intranuclear puncta; black arrows mark examples of cytoplasmic puncta. Scale bar, 5 µm, for all images, except for 2.5-fold enlarged images in inset. For germline location, see **Figure S5A**.

**Table S1.**
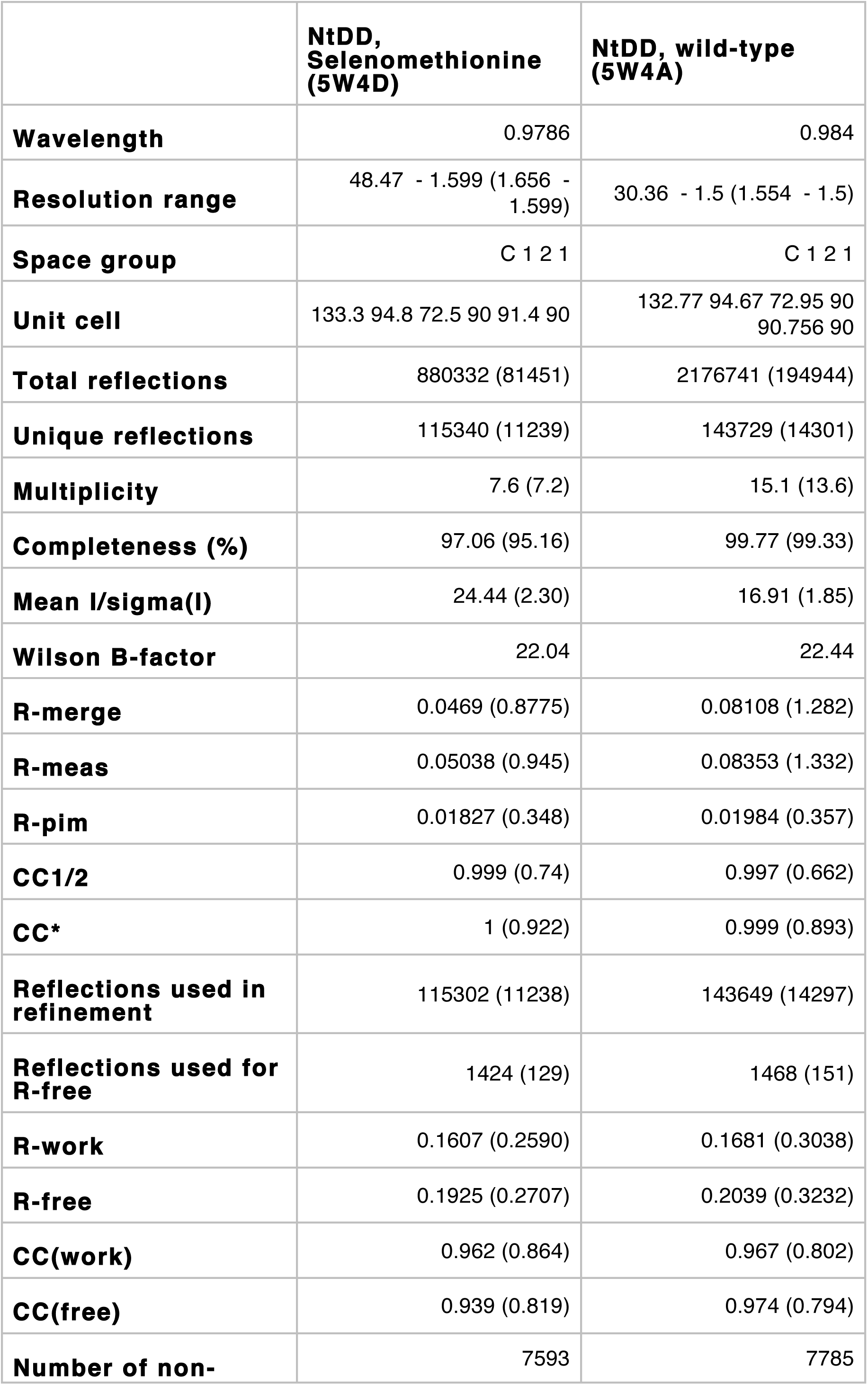
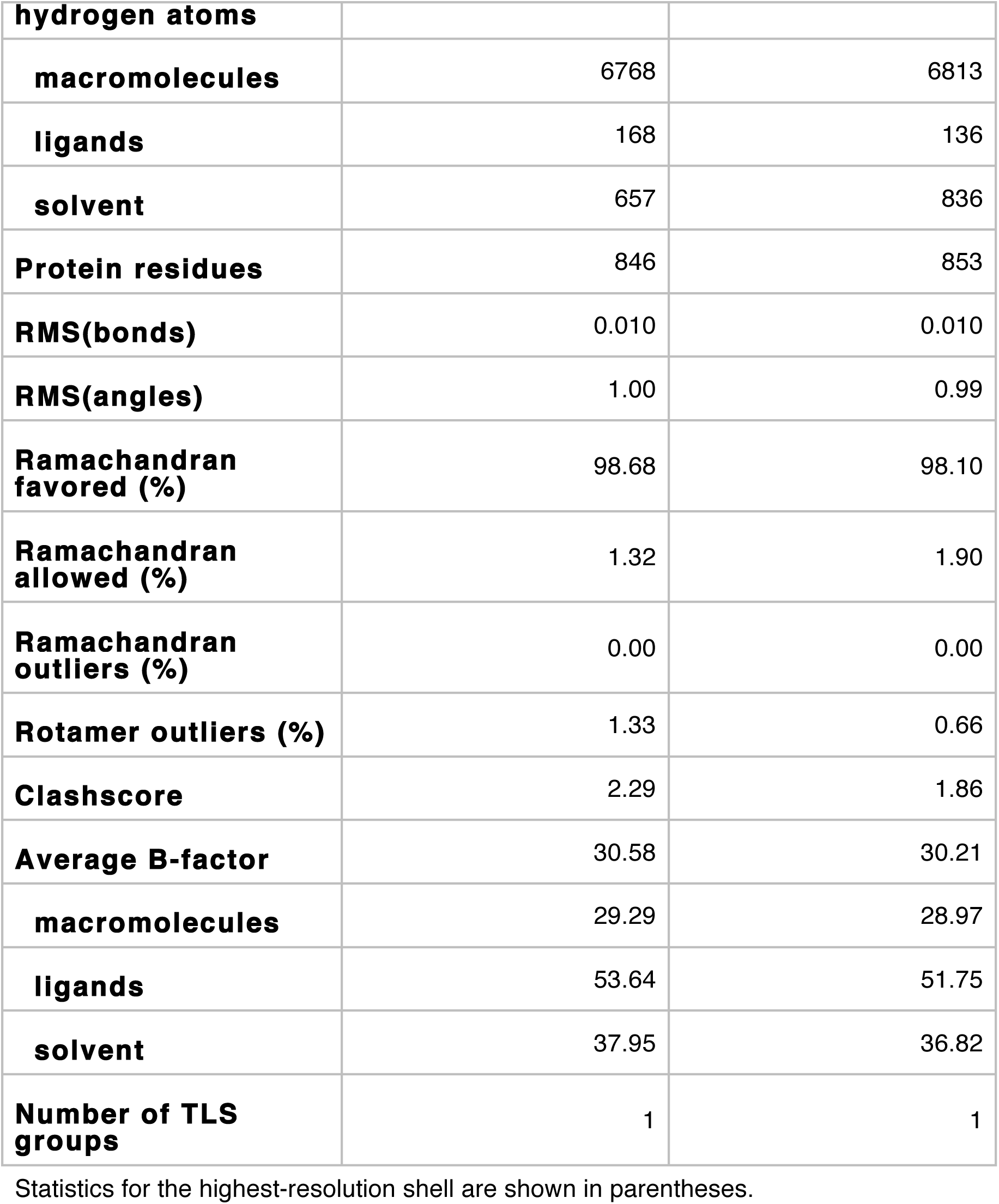
*C. japonica* PGL-1 NtDD crystal structure data and model statistics

